# Plasma-derived exosomal analysis and deconvolution enables prediction and tracking of melanoma checkpoint blockade response

**DOI:** 10.1101/809699

**Authors:** Alvin Shi, Gyulnara G. Kasumova, William A. Michaud, Jessica Cintolo-Gonzales, Marta Díaz Martínez, Jacqueline Ohmura, Arnav Mehta, Isabel Chien, Dennie T. Frederick, Sonia Cohen, Deborah Plana, Douglas Johnson, Keith T. Flaherty, Ryan J. Sullivan, Manolis Kellis, Genevieve M. Boland

## Abstract

**Purpose:** Immune checkpoint inhibitors (ICI) have demonstrated promising therapeutic benefit although a majority will not respond. Here we identify and validate predictive biomarkers from plasma-derived exosomes that allow non-invasive monitoring of tumor intrinsic and host immune status and prediction of ICI success.

**Experimental Design:** Transcriptomic profiling of peripheral blood bulk exosomes and tumors from a discovery cohort of 50 patients with metastatic melanoma treated with ICI was undertaken; a further validation cohort of 30 patients was utilized to validate findings from the discovery cohort. We designed a Bayesian probabilistic model to partition bulk exosomes into tumor-specific and non-tumor-specific proportions.

**Results:** Exosomal RNA signatures exhibit significant correlations with tumor transcriptomes. Exosomal profiles reflect several key biological drivers of ICI resistance or melanoma progression, exhibit significantly differentially expressed genes and pathways, and correlate with and are predictive of clinical response to therapy. Our deconvolution model estimates contributions from tumor and non-tumor sources, enabling more precise interpretation of differentially-expressed genes and pathways. Exosomal RNA-seq mutational information can be used to segregate responders and non-responders.

**Conclusions:** Peripheral blood-derived exosomes can serve as a non-invasive biomarker to jointly probe tumor-intrinsic and immune changes to ICI, and can potentially function as predictive markers of ICI responsiveness and a monitoring tool for tumor persistence and immune activation.

**Statement of Significance:** We use transcriptomic analysis of bulk, non-selected, peripheral blood derived exosomes to reveal both tumor-intrinsic and immune-derived signatures predictive of early response to immune checkpoint inhibitor therapy. We develop a novel computational model to classify exosomal transcripts into tumor and non-tumor components and establish relevance in immune checkpoint blockade therapy. We show that tumor driver load from RNA-seq mutational calls are significantly different between responders and non-responders.

## Introduction

Currently available blood-based biomarkers for immunotherapy resistance focus on cell-free DNA (cfDNA) or circulating tumor cells^1, 2^, which solely reflect tumor-based properties and do not provide insights into the underlying transcriptomic changes occurring in the immune system during ICI treatment. To improve prediction and tracking of ICI resistance, simultaneous capture of transcriptomic features from both the tumor and immune system^3^ is critical.

Exosomes are circulating extracellular vesicles (EVs) that contain a subtranscriptome of their cell of origin and are produced by many cell types, including tumor and immune cells. Exosomes are involved in oncogenesis, immune modulation, and serve as communicators of genetic and epigenetic signals. In several cancers tumor-derived exosomes (TEX) can modulate the tumor microenvironment and elicit anti-tumoral immune responses^4^, and plasma-derived exosomal transcripts are markers of anti-tumor immune activity^5^. Exosomes are secreted by many immune cell-types implicated in ICI response including CD4+ and CD8+ T-cells, dendritic cells, regulatory T-cells, and macrophages^6–9^. Work utilizing exosomes for cancer diagnostics enrich for TEX and exclude exosomes from other sources^10^. We hypothesize that isolating bulk, non-enriched exosomes captures both tumor-derived and non-tumor-derived exosomes reflect both tumor-intrinsic and non-tumor-intrinsic mechanisms of ICI. We analyzed pre-treatment and on-treatment peripheral blood-derived bulk exosomal RNA from a discovery cohort of 50 patients with metastatic melanoma treated with ICI via transcriptome microarray. Our discovery cohort consisted of 33 responders and 17 non-responders (Fig. S1a and **Table S1**). A subset of patients had post-treatment plasma samples (N=15) and tumors (N=26). Additionally, we profiled four melanoma cell lines and their paired exosomes. We validated our discovery cohort findings in a validation cohort of 30 patients (N=19 responders, N=11 non-responders) using exoRNA-seq. To deal with transcriptomic data from two different sequencing platforms, we utilized a multi-pronged statistical analysis strategy in order to minimize cross-platform variance for relevant analyses (Fig. S1b, **Methods**) To interpret the mixed plasma-derived exosomal data, we developed a novel Bayesian deconvolution computational model to partition exosomal transcripts into tumor and non-tumoral sources.

## Results

To correlate exosomal transcriptomes with tumor transcriptomes, we examined the relationship in expression between melanoma cell-lines and cell-line-derived exosomes. We observed high levels of correlation between cell-lines and their exosomes (average R^2^=0.87, Fig. S3a). Cell-lines shared similar concordance in gene expression (Fig. S3b), and the majority of genes had small differences in overall expression (Fig. S3c). Unsurprisingly, each cell line had the highest correlation with their corresponding exosomes and not exosomes from other cell lines (Fig. S3d). This suggests that exosomes are reasonable proxies for melanoma expression in patient derived cell-lines. Next, to determine if patient plasma-derived exosomal transcripts correlated with patient tumors, we analyzed paired exosomal and tumor transcriptomes from N=9 patients and saw close correlation of expression (average R^2^=0.82, Fig. 1a). Concordance analysis demonstrated that most genes expressed in tumors are also detected in corresponding exosomes (Fig. 1b). By analyzing genes unique to tumors versus exosomes, we found enrichment of immune-related signatures exclusively in exosomes (e.g., T-cell activation, NK activation), while tumor-exclusive transcripts enriched for metabolic and tumor-related pathways (Fig. 1c, **Table S2**). To characterize cell populations present in exosomes and whether they corresponded to populations present in the tumor microenvironment, we utilized CIBERSORT to computationally infer immune cell-type enrichments in our patient plasma-derived exosomal samples^11^. We observed a relative enrichment in 5 immune sub-populations exclusively in exosomes, including neutrophils, natural killer cells, and CD4+/CD8+ T-cells, and a relative depletion in macrophages and mast cells (Fig. 1d). This suggests that exosomes are over-enriched for signals from specific immune populations that play key roles in anti-PD1 responses^3^. Based on these results, we hypothesized that plasma-derived exosomal transcripts from patients prior to and during treatment would predict or reflect ongoing resistance to ICI.

**Figure 1:**
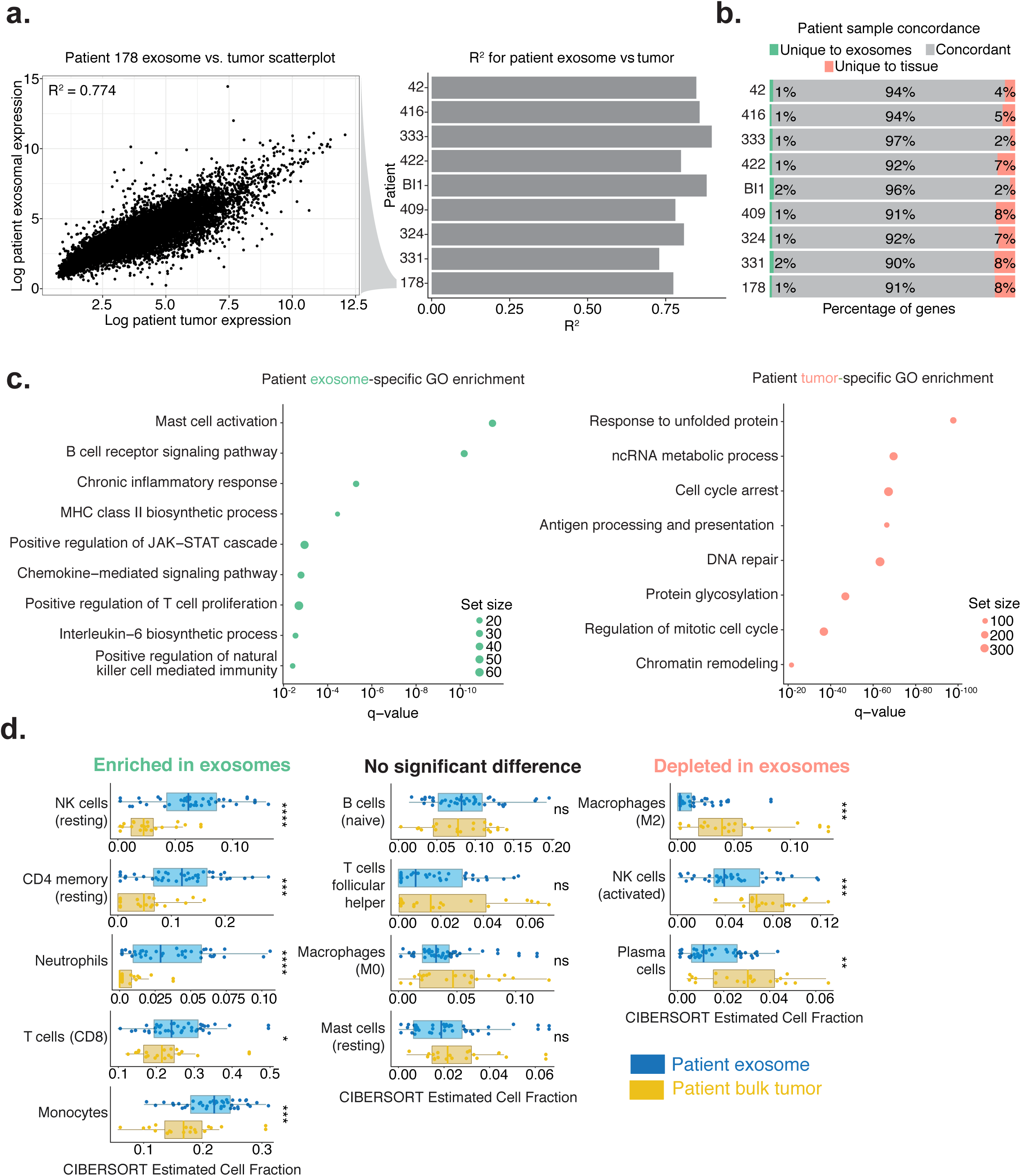
Tumor and exosomal RNA concordance. Characterization of transcriptomic similarities between patient melanomas and time-matched exosomes. (a) Scatter plot displaying the relationship between expression values of tumors and plasma exosomes in a representative patient (Patient 178) and a histogram R^2^ across the cohort. If a patient had multiple samples of the same type at the same time-point, the samples were averaged prior to computing the R^2^. (b) Concordance was calculated using a low-expression threshold cut-off for expressed versus non-expressed status (methods). Genes expressed or not expressed in tissue and exosomes were considered concordant (blue), while a subset of transcripts were unique exosomes (red) or tumor (green). (c) Enrichment of genes unique to patient exosomes (left) or patient tumors (right) and GO enrichment analysis on unique genes from the categories in panel b. (d) CIBERSORT inferred deconvolution estimates for all pre-treatment patient tumor and pre-treatment patient plasma-derived exosomal samples using LM22 immune reference profiles. Technical replicates were averaged and biological replicates were considered independently. The data is segregated into three categories based on the results of a Mann-Whitney U-Test between exosomal and tumor inferred CIBERSORT fractions for each cell-type.

Since tumoral post-treatment signatures are more representative of ICI response than pre-treatment values^1, 12^, we assessed the ability of on-treatment exosomal transcripts to reflect ongoing ICI responses in both a validation and discovery cohort. Through differential gene set analysis utilizing the canonical pathways gene set from the Molecular Signatures Database (MSigDB)^13^ via Gene Set Variation Analysis (GSVA, Methods)^14^, we observed 258 pathways with significant differences between responders and non-responders in our discovery cohort, of which 25 pathways were also significant in our validation cohort. Many validated pathways, such as T-cell receptor, CTLA4, TGF-β, SMAD2/3, Notch, TNFR2, and VEGFR signaling pathways (Fig. 2a, SFig. 4), are related to previously implicated mechanisms of ICI resistance or melanoma progression and metastasis^15–18^. Since our longitudinal dataset includes multiple on-treatment time points, we are able to visualize differential on-treatment pathway dynamics via single-sample GSVA scores^14^. We observed a noticeable degradation of T-cell receptor (TCR) pathway activity during the course of treatment in non-responders, suggesting that ICI treatment failed to restore T-cell activity in non-responders (Fig. 2c, SFig. 5a). This is further supported by visualizing the dynamics of CD28 co-stimulatory pathway, which mirrors the decline in TCR signaling (SFig. 6a). We also observe divergence in CTLA4 pathway activity levels over time (SFig. 6b), potentially as a result of peripheral tolerance in response to ICI treatment^19^. Furthermore, we also observe temporal divergence in tumor-related pathways, such as P53-Hypoxia (SFig. 6c) and Kinesin activity (SFig. 6d). At the individual gene level, there were 1240 differentially expressed genes (DEGs) in the discovery cohort and 514 DEGs in the replication cohort. 80 genes shared successful p-value validation and 47 were successfully replicated at both the p-value, minimum expression, and log-fold change levels (Fig. 2c, **Table S3**). The 47 replicated DEGs represents a replication rate significantly above that expected solely by chance (p=0.00088, hypergeometric test). Within the replicated DEGs, we observed a number of DEGs that mirrored our findings from our gene-set enrichment analysis. For example, KLF10 is a major effector in the TGF-β signaling pathway^20^ and WNT8B’s role in Wnt signaling may impact T effector cell differentiation^21^. Moreover, we observed the presence of several melanoma related antigens (MAGEA1, MAGEA3) in our validated DEGs that are known cancer testis antigens recognized by cytolytic T-cells in melanoma and are almost exclusively expressed by melanoma cells^22^. To better illustrate the time dynamics of our on-treatment DEGs, we plotted the normalized expression changes relative to the first collection time (Fig. 2d) and the unnormalized expression changes (Fig. S5b-e) for several validated DEGs (MAGEA1, MAGEA2, KLF10, and MIR4519). These plots demonstrate that the on-treatment DEGs such as MAGEA1, KLF10, and MIR4519 also have differential ICI-induced dynamics between responders and non-responders in their transition from pre-treatment to on-treatment time points.

**Figure 2:**
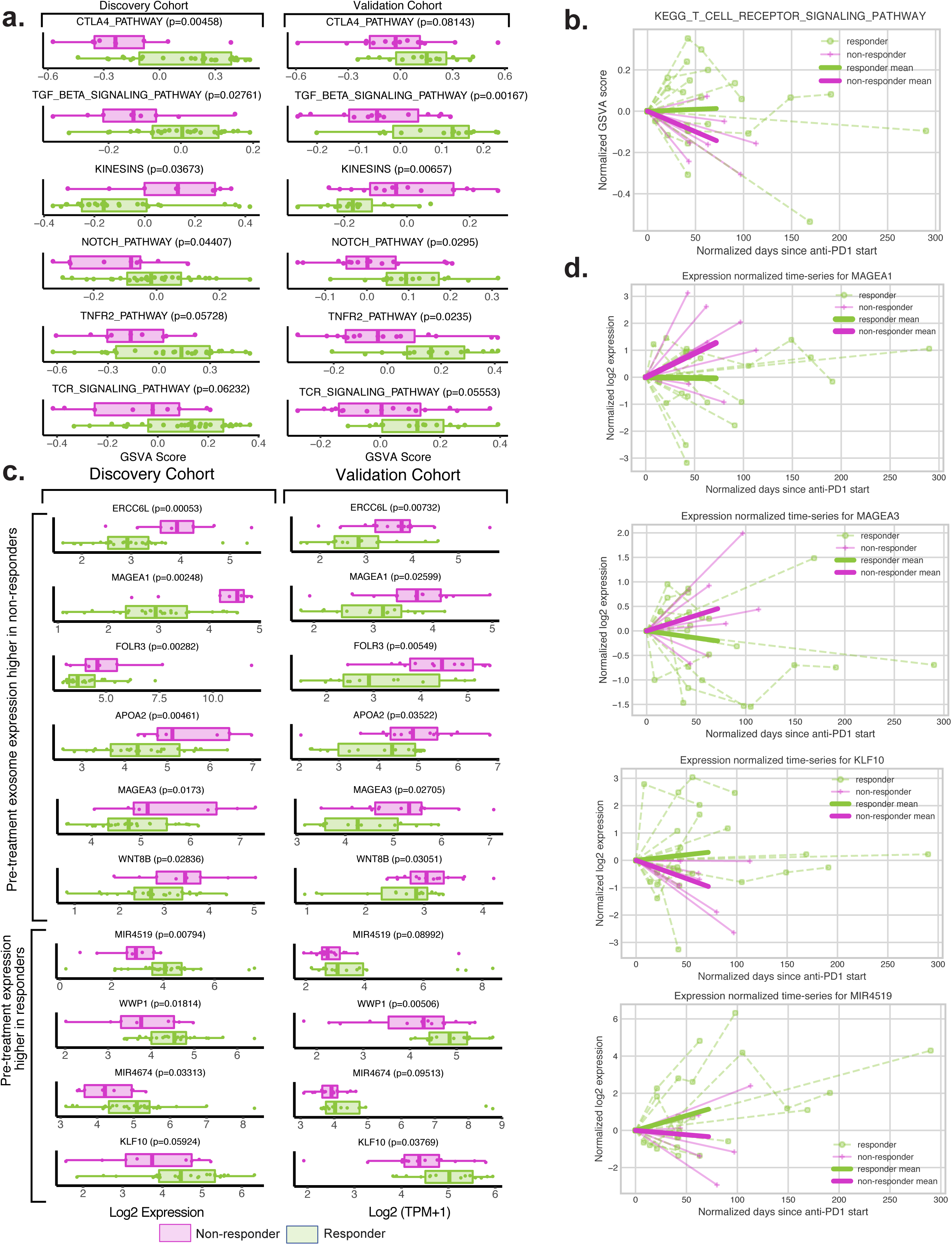
Biological pathways and genes that stratify responders and non-responders in on-treatment exosomes. (a) MSigDB canonical (C2) pathway enrichments in on-treatment samples, (salmon color: discovery cohort; teal color: validation cohort). (b) Time dynamics of T-cell receptor KEGG signaling in responders versus non-responders visualized using single-sample gene set enrichment scores derived from GSVA analysis. Individual patient progressions are displayed as connected lines. Both the start time and the relative expression levels were normalized against the time and GSVA score at the first sample collection point, which were set to 0. The mean GSVA score changes for responders and non-responders are plotted in bold green (responders) and purple (non-responders) lines. (c) Expression comparison between responders (green) and non-responders (purple) for selected validated differentially expressed genes between responders (green) in on-treatment samples across both the discovery and validation cohort. The p-values displayed are generated from limma. (d) Time dynamics of several representative validated on-treatment DEGs, including MAGEA1, MAGEA3, KLF10, and MIR4519 in the discovery cohort. Individual patient progressions are displayed as connected lines. Both the start time and the relative expression levels were normalized against the time and expression at the first sample collection point, which were set to 0. The mean expression changes for responders and non-responders are plotted in bold green (responders) and purple (non-responders) lines.

We next assessed if pre-treatment exosomal transcriptomes are able to stratify responders from non-responders and reflect mechanisms underlying ICI resistance. We performed gene-set enrichment of pre-treatment responders vs. nonresponders via GSVA^23^, which showed 101 differentially expressed MSigDB canonical pathways in the discovery cohort, of which 26 replicated in the validation cohort (Fig. 3a, SFig. 7)^24^. Comparing the on-treatment pathways, we see that differential Notch and TGF-β signaling is also found in the pre-treatment cohort. We observe differences in MAPK related signaling (ERRB4) and pathways reflective of melanocytic processes (keratinization). We found 366 and 1406 differentially expressed genes (DEGs) at the pre-treatment time point in the discovery cohort and validation cohort, respectively, of those, 54 DEGs had replicated p-values while 38 genes had replicated both p-values and log fold changes. This represented a replication rate significantly above that of random chance (p=0.0041, Hypergeometric test). Our DEGs included members of both immune and tumor-related pathways implicated in ICI resistance or tumor growth in our pathway level analysis, such as CD1A, MAP2K4, TRBV7-2 and IFGL1^15, 25, 26^ (Fig. 3b, **Table S4**). Cancer-associated miRNAs, such as miR551A, were also enriched in nonresponders. To assess the predictive ability of the replicated DEGs, we constructed a pre-treatment random forest classifier that achieved high predictive performance (peak AUC=0.805) in the discovery cohort using k=5 stratified cross-validation using the validated pre-treatment DEGs (Fig. 3c). To independently confirm the predictive power of our model, we utilized a 10-tree random forest classifier with trained on the discovery cohort to test in our independently generated validation cohort. We found that our model is moderately predictive in a novel cohort (AUC=0.84, Fig. 3c). We also found that multiple validated DEGs, specifically IGFL1, TFF2, and MAP2K4, showed significant stratification with regards to either progression-free or overall survival (Fig. 3d, SFig. 8).

**Figure 3:**
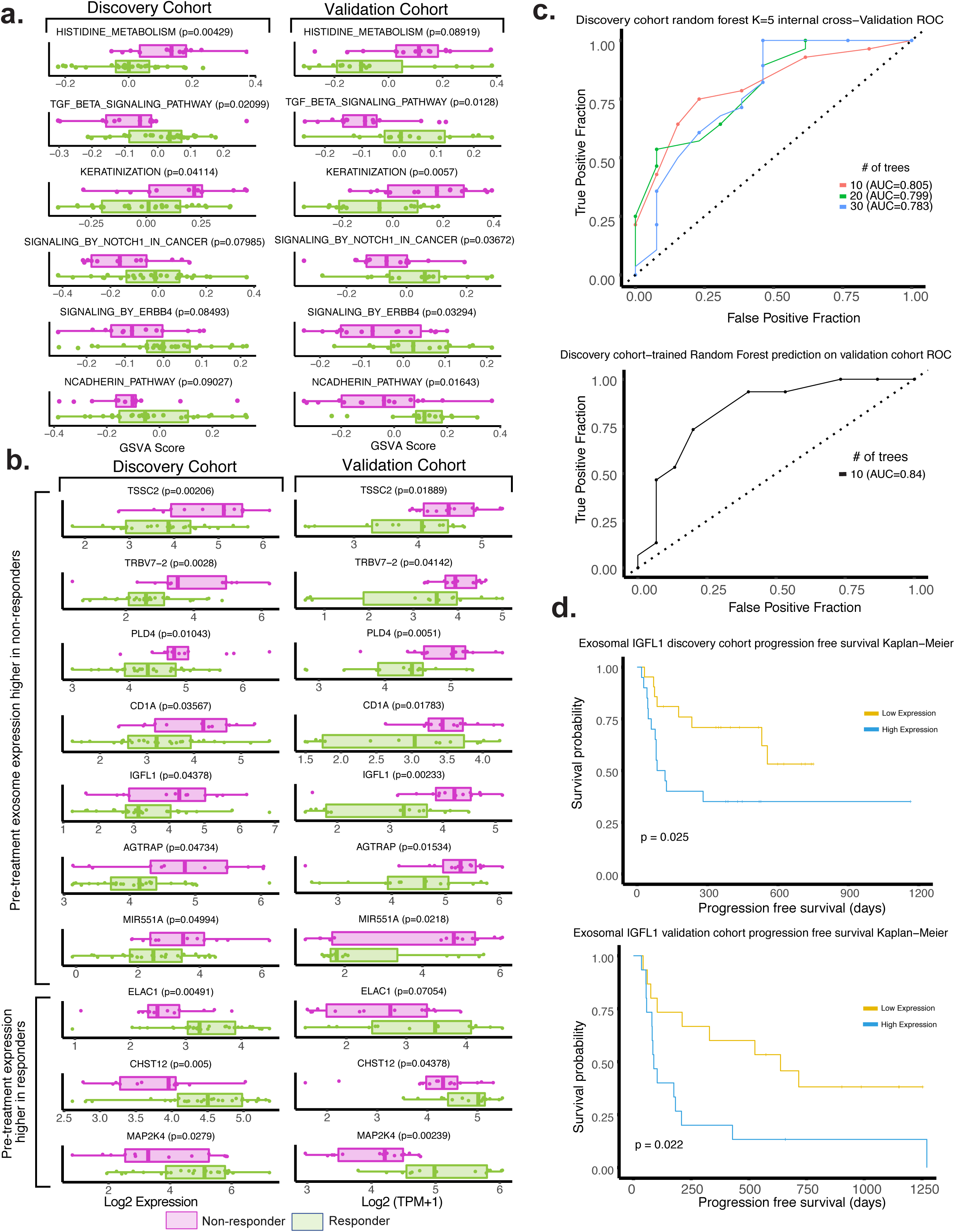
Biological pathways and genes that stratify responders and non-responders in pre-treatment exosomes. (a) GSVA scores and associated p-values for selected MSigDB canonical pathways that different significantly between responders (green) and non-responders (purple) in pre-treatment exosomes. The p-values were generated by performing a Mann-Whitney U-Test between responder and non-responder GSVA scores. (b) Boxplots of expression values of selected validated pre-treatment exosomal DEGs between responders and non-responders in both the discovery and validation cohort (purple color: non-responders, green color: responders). (c) Receiver operating characteristics using a random forest classifier with the validated DEGs genes between responders and non-responders as features. We tested 10, 20, and 30 trees random forests using default parameters from the python package ‘sklearn’^31^. We used the best performing model - random-forest with 10 trees - on the validation cohort to obtain out-of-sample prediction performance. (d) Kaplan-Meier overall survival plots for IGFL1 in the discovery and validation cohorts.

The non-selected bulk exosome approach raises questions regarding how exosomes from non-tumor sources impact estimations of tumor-derived exosome contributions. Based upon CIBERSORT results (Fig. 1c) we hypothesized that we detect both tumor and non-tumor exosomes. Understanding the contribution from tumor-derived versus non-tumor derived exosomal sources may define whether changes in plasma-derived exosomes during ICI treatment reflect changes in the tumor microenvironment or non-tumoral changes (i.e. systemic changes in the immune system). Therefore, we developed a novel probabilistic deconvolution model to infer: (i) a “packaging” coefficient that represents the preferential depletion/enrichment of transcripts during packaging and export into exosomes, (ii) the unobserved non-tumor derived exosomes (SFig. 9), and (iii) a mixing fraction between unobserved tumor-derived exosomes and non-tumor-derived components for each gene (Fig. 4a). To validate our model, we benchmarked our deconvolution findings with a bulk *in silico* deconvolution algorithm CIBERSORTx^27^ and found overall concordance between deconvolution predictions and those generated using an independent algorithm and reference profiles (**Supplemental Note**). Our deconvolution model addresses the relative contributions of bulk tumor and non-tumor sources in DEGs and key genes involved in anti-PD1 response. Analysis of known genes allowed us to test the accuracy of the deconvolution predictions in a known gene set (Fig. 4b), followed by validated DEGs at both the pre and on treatment timepoints (Fig. 4c). Our model can also probe relative enrichment across gene sets and calculate a gene-set level tumor fraction. We can also visualize the global landscape of inferred packaging coefficients (Fig. 4d), demonstrating significant enrichment for tumor-derived transcripts. When visualizing a number of ICI and melanoma-relevant KEGG pathways, our results also align with expected ranking (e.g. melanoma-related pathways have higher tumor fraction) (Fig. 4e). To examine whether our pre-treatment and on-treatment validated DEGs preferentially enrich for tumor-derived or non-tumor-derived genes, we visualized the fraction of genes in each group predicted to be derived from non-tumor sources. We see that pre-treatment validated DEGs are preferentially depleted for non-tumor genes (and enriched for tumor-derived genes), whereas on-treatment validated DEGs have greater non-tumor contribution (Fig. 4f). This suggests that on-treatment DEGs preferentially reflect immune-related changes induced by ICI treatment, consistent with our findings from our on-treatment differential pathway analysis. To illustrate the utility of our deconvolution algorithm as a mechanism for interpreting our DEGs, we use KLF10, a member of the TGF-β pathway that has both roles a tumor suppressor^20^ and plays a role in inducing Th1/Th17 polarity and CD8+ memory T-cell formation^28^. Both roles may reflect plausible mechanisms of resistance to anti-PD1 therapy; however, our data shows that KLF10 is mostly derived from the non-tumor component, suggesting that the differential on-treatment response may be correlated with potentially with the Th1/Th17 polarity inducing function of KLF10 as opposed to its tumor suppressor role in mediating resistance to anti-PD1 therapy.

**Figure 4:**
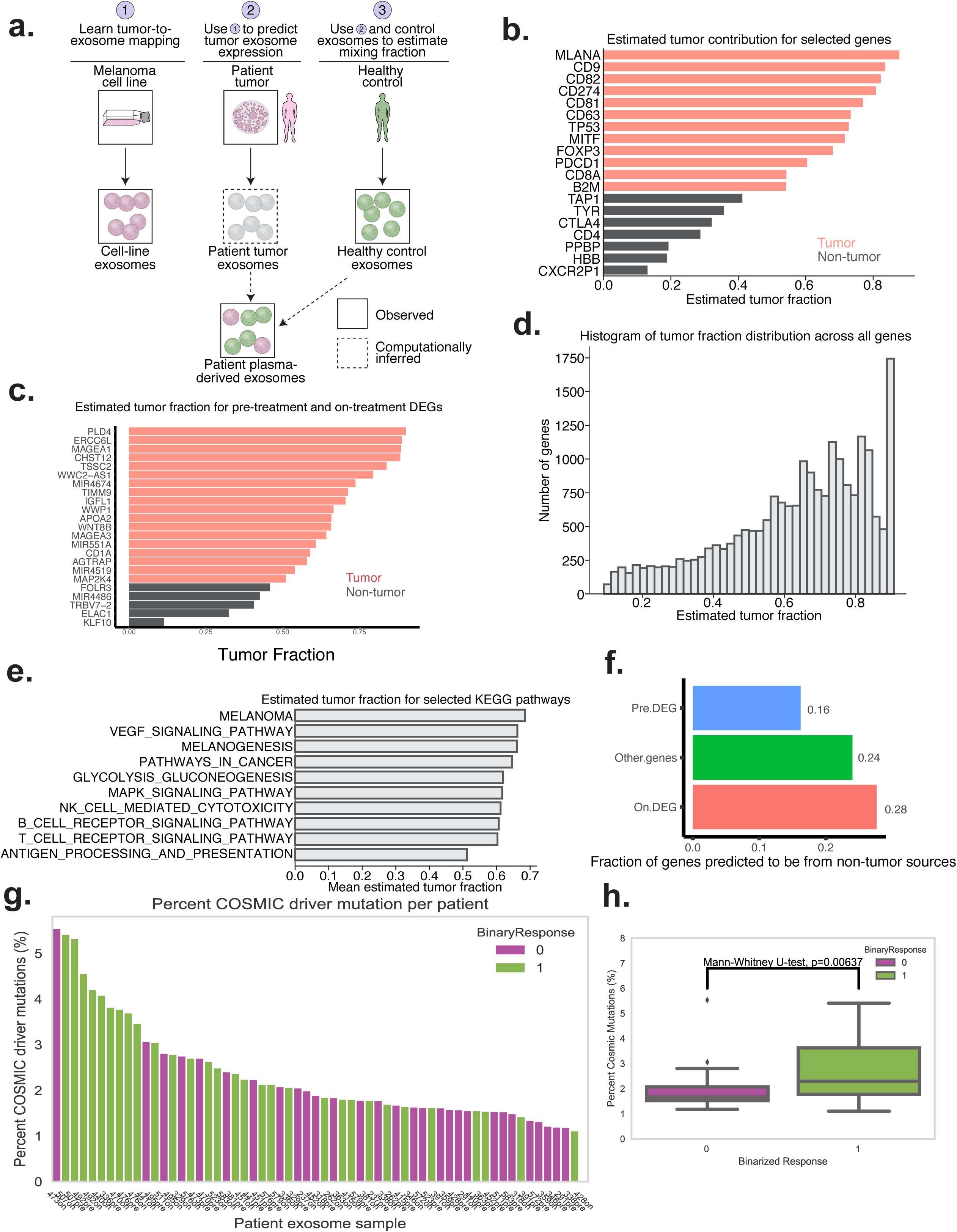
Deconvolution of exosomal profiles and analysis of driver mutations in RNA-seq profiles. (a) Schematic representation of our deconvolution model (see Supplementary Note). (b) Selected tumor contributions for known tumor and non-tumor genes. Red denotes predicted tumor and grey denotes predicted non-tumor. (c) Estimated tumor fraction for pre-treatment and on-treatment DEGs demonstrating that most DEGs higher in non-responders are predicted to come from non-tumor sources, while DEGs enriched in nonresponders (miRNA) are predicted to derive from tumors (d) Histograms of maximum a posterori (MAP) estimates of tumor fraction from our model across all genes. (e) Average tumor fraction of all genes involved in several selected KEGG categories. (f) Fraction of validated pre-treatment DEGs, on-treatment DEGs, and all other genes predicted to be non-tumor derived (i.e., predicted tumor fractions of <0.5) (g) Percentage of COSMIC driver mutations as a part of the total mutations (a combination of somatic & germline) called for each patient’s RNA-seq profile in the validation cohort showing a higher mutational load in responder patients (purple color: non-responders, green color: responders). (h) Boxplot summarizing the distribution of COSMIC driver mutation fraction between responder and non-responder profiles. A Mann-Whitney U-test was used to test for significant differences between the responder and non-responder distributions.

Finally, we investigated whether mutational information embedded in the exoRNA-seq data can be utilized to stratify responders and non-responders, due to the success of tumor mutational burden (TMB) via tumor WES and cfDNA as an indicator of tumor immunogenicity and as an ICI biomarker^29^. Despite not having matched normal WES data for our validation cohort, we were able to survey the mutational landscape by using hg38 genomic reference to call both somatic and germline RNA-seq-associated mutations against using our exosomal RNA-seq data (methods). By comparing our exoRNA-seq mutational calls against patient matched tumor panel sequencing results, we determined that three of our patients had specific driver mutations called by both in-house NGS panel sequencing and our RNA-seq mutational calling pipeline (**Table S1**). Panel sequencing data represents only a small fraction of somatic tumor-associated mutations; to gain a more comprehensive view, we also surveyed the entire somatic mutational landscape to determine whether there are significant differences in cancer driver mutational burdens between responders and non-responders. Although the patient samples varied in the absolute number of mutations detected, we reasoned that significant differences in the fraction of somatic tumor-related mutations from the COSMIC database relative to the overall mutational pool can be largely attributed to changes in the somatic mutation load present in the exosomal RNA and not significant differences in germline mutations^30^. In concordance with previous observations from WES and cfDNA TMB estimates, we observed that responders tended to have significantly higher COSMIC driver somatic mutational fraction relative to non-responders (Fig. 4g-h).

## Discussion

In this study, we explored the potential usage of plasma-derived exosomal transcriptomic profiles as a source of biomarkers for predicting and monitoring checkpoint blockade immunotherapy success. Our results show that exosomes, in aggregate, correlate with certain aspects of bulk tumoral biology and reflect a number of previously identified differentially-regulated biological pathways implicated in ICI resistance or melanoma progression in responders vs. non-responders. The majority of differential pathways and genes at the pre-treatment time point primarily reflect differences in metabolic state as opposed to pre-existing immune-related differences, suggesting that plasma-derived exosomes only start capturing immune-related differences between responders and nonresponders after ICI treatment is administered. This is supported by the enrichment of immune-related pathways in our on-treatment differential pathway analysis, as well as the enrichment for non-tumor-derived DEGs as inferred by our deconvolution model. Though the validated DEGs we discovered are biologically informative and can be utilized to create predictive models, the uncertain tissue-of-origin of exosomal transcripts confounds our ability to define a mechanistic basis for differential expression. To address this, we developed a novel Bayesian deconvolution model uniquely suited to deconvoluting plasma-derived exosomal transcriptomic expression. To our knowledge, this is the first computational model that explicitly models preferential packaging of exosomes and contribution of non-tumor exosomal components in order to infer per-gene tumor fractions.

The higher levels of correlation between melanoma cell-lines and their exosomes as compared to bulk patient tumors and corresponding plasma-derived exosomes suggest that bulk plasma-derived exosomes reflect a more broad repertoire of exosome sources. Indeed, the most robust enrichment in exosomes is for immune-related pathways. This is reinforced by the relative enrichment of several key immune cell-types in our CIBERSORT deconvolution and are validated by our on-treatment DEGs and pathway enrichments. Exosome transcriptomic biomarkers may complement circulating tumor DNA (ctDNA) to gain transcriptomic information regarding tumor dynamics in addition to genomic information and may give a readout of both tumor-intrinsic and immunologic changes simultaneously.

To address tissue of origin of exosomal transcripts, we developed a deconvolution model to characterize exosomal transcripts from tumor versus non-tumoral sources that explicitly accounts for differential exosomal transcript packaging. Currently, our model can only differentiate between tumor versus non-tumor contributions; however, ongoing experiments utilizing cell-specific exosome selection and/or depletion may enable us to differentiate between specific sources. Our deconvolution model is limited by three major factors: (i) the simplifying assumptions regarding the linear nature of the packaging coefficient and how its shared between *in vitro* and *in vivo* samples; (ii) lack of accounting for both tumor and patient heterogeneity, which is in part due to (iii) the limited number of samples. As we continue to analyze data from more patients and perform *in vitro* exosome selection experiments, we anticipate that our current model will serve as the foundation for more sophisticated models that can address these issues. Despite these limitations, our deconvolution model is able to pinpoint the potential source of exosomal expression and generate testable hypotheses. Work is ongoing to experimentally validate the predicted source of circulating exosomes via both tumor and immune cell-specific exosome selection, which will be used to iteratively improve our deconvolution model and establish potential causal roles for exosomal transcripts in driving ICI resistance.

Our utilization of exoRNA-seq technology in the validation cohort brought additional challenges to our analysis when cross-comparing with discovery cohort microarray data. We reason that utilizing two separate sequencing technologies on two independent cohorts raises the bar for reproducibility and that findings replicated with distinct methodologies are likely to be robust. Additionally, we show that mutational information embedded in the exoRNA-seq itself can potentially be exploited to stratify responder and non-responder populations and serves as an orthogonal means to determine the tissue-of-origin of plasma-derived exosomal transcripts. This approach can be further enhanced through complementary WES of exosomal DNA. Although it is unlikely the utility of the mutational information from exosomal RNA-seq data will outstrip high-depth cfDNA or WES TMB data in the near future, this mutational information is embedded within a large amount of transcriptomic information provided which reflects dynamic tumoral changes and can complement other DNA-based sequencing methods (ctDNA, exoDNA-seq) in ICI monitoring and response prediction tasks.

## Supplementary figures

**Supplementary figure 1:**
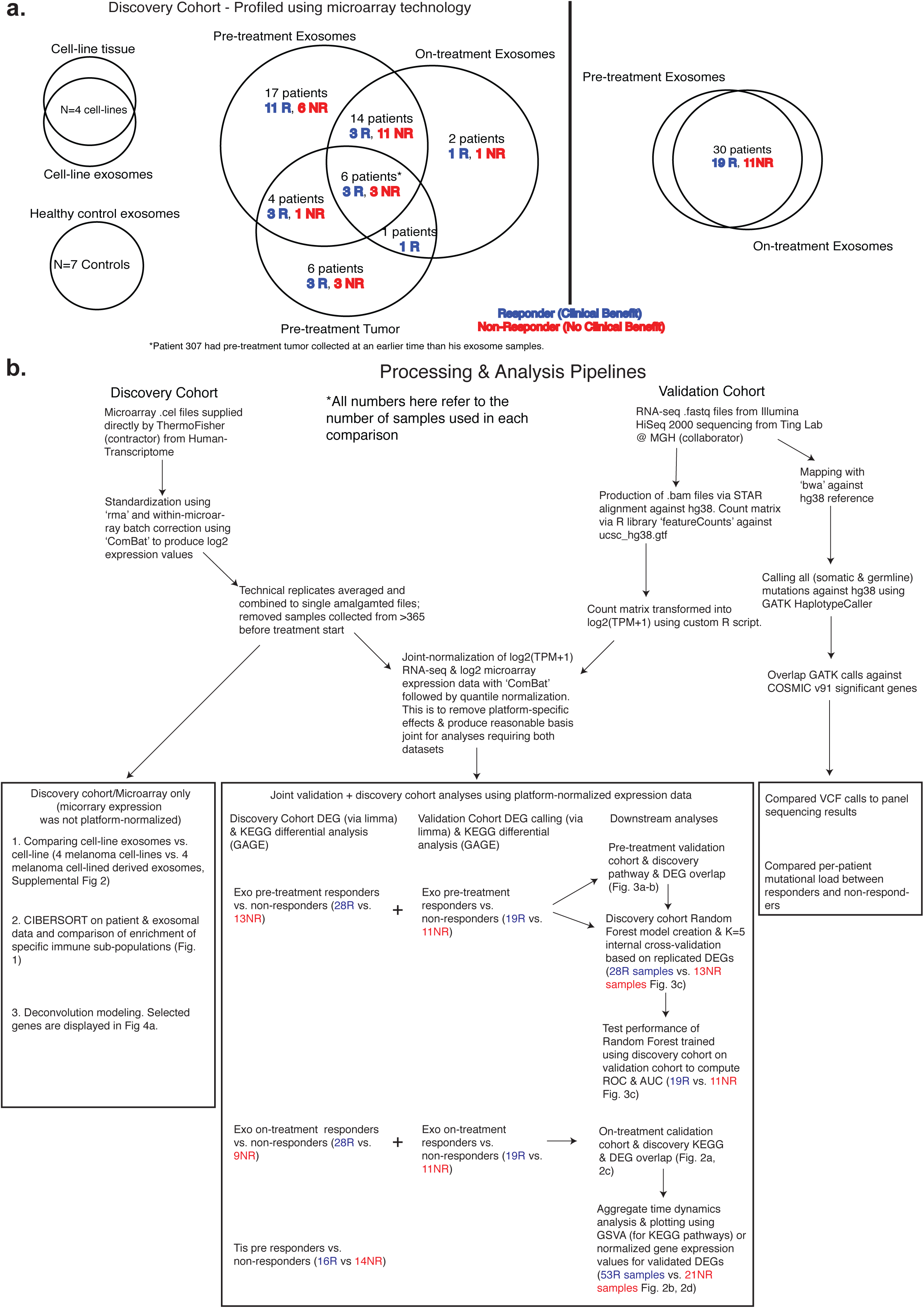
(a) Outline of both the discovery and validation study cohort. The complete cohort metadata is available in **Supplementary Table 1**. (b) Outline of the processing and analysis steps undertaken to generate the major results in the paper.

**Supplementary figure 2:**
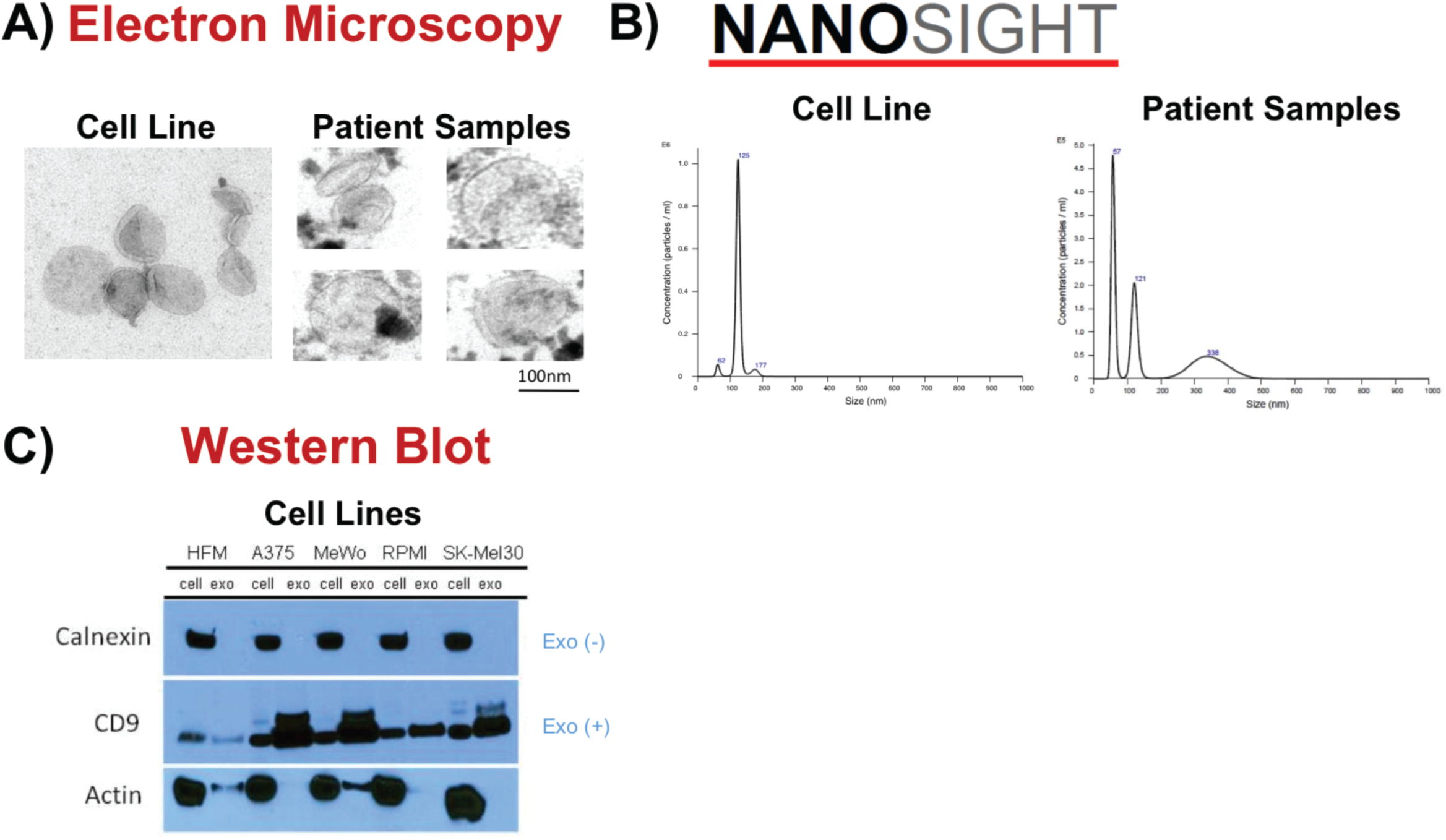
In vitro exosome characterization. (a) Electron microscope images of representative cell-line and patient-derived exosomes. (b) Nanosight analysis of a representative cell line sample (RPMI) and patient sample. (c) Western blot of 5 cell line/paired EV protein with no calnexin within exosomes, but high levels of CD9 within exosomes as compared to paired cells.

**Supplementary figure 3:**
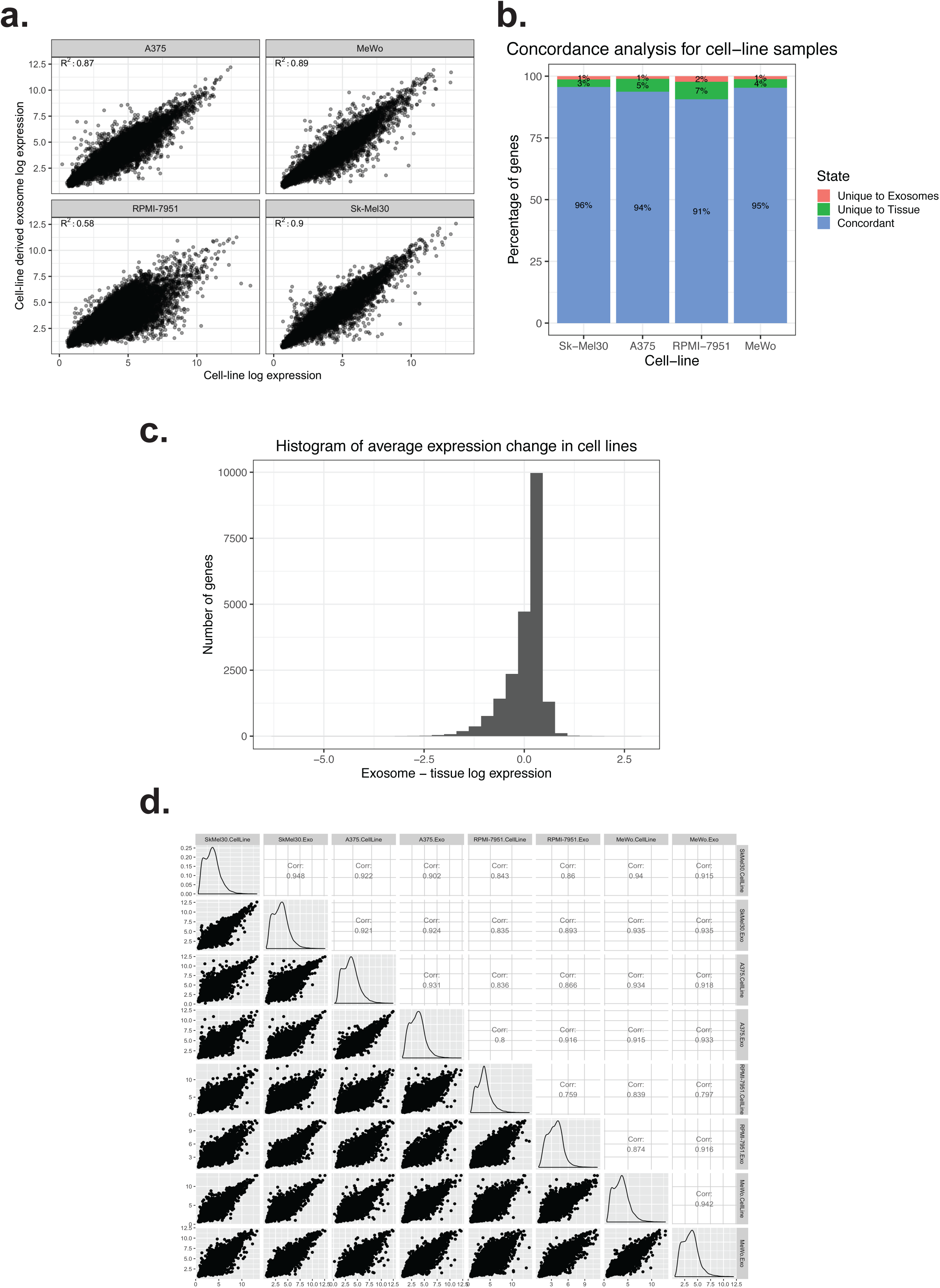
Characterization of transcriptomic similarities between cell-line derived melanoma samples and their matched exosomes. (a) Scatter plot visualizing differences between tissue and plasma-derived exosomes in patient samples. (b) Histogram visualizing the log fold changes between melanoma cell-lines and their exosomal counterparts. The profiles were compiled using the average of four expression profiles. (c) Concordance analysis across our 4 cell-line samples. Concordance was calculated by using a low expression cutoff as a cut-off for expressed vs. non-expressed status. Genes that were either expressed or not expressed in both tissue and exosome compartments are considered concordant. (d) Correlation plot between all 4 cell-lines and their exosomal counterparts.

**Supplementary figure 4:**
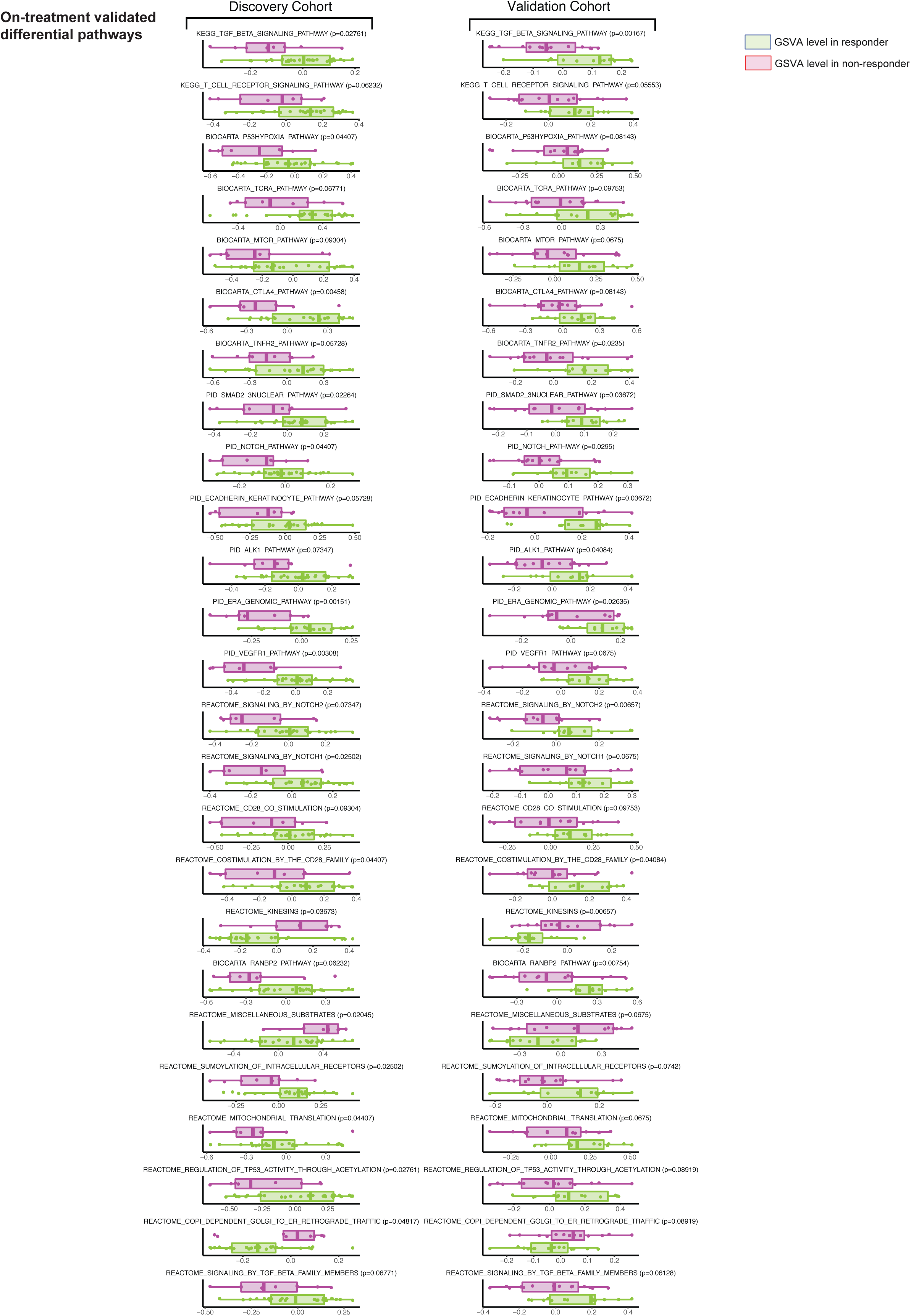
Box-plots and associated p-values for validated MSigDB canonical pathways that differ between responders (green) and non-responders (purple). The visualized points are individual GSVA scores inferred for each pathway. The p-values were generated by comparing responder vs. on-responder GSVA scores via a Mann-Whitney U-test.

**Supplementary figure 5:**
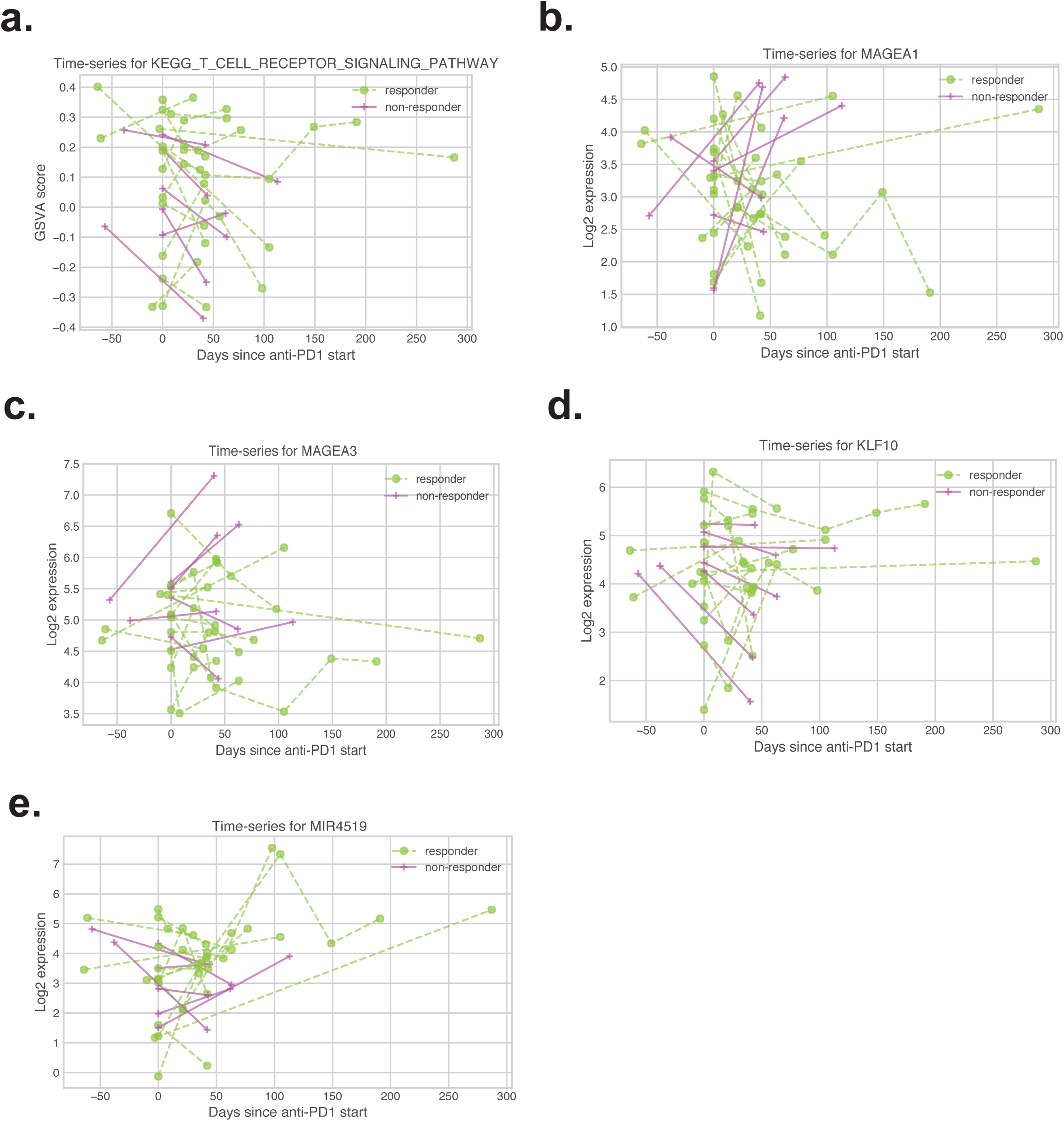
(a-e) Un-normalized time-series plots showing time dynamics for pathways and genes discussed in Fig. 2b and Fig. 2d. Individual GSVA scores were used to plot the TCR KEGG pathway, while platform-normalized log2 expression values were used to plot the individual gene expression levels.

**Supplementary figure 6:**
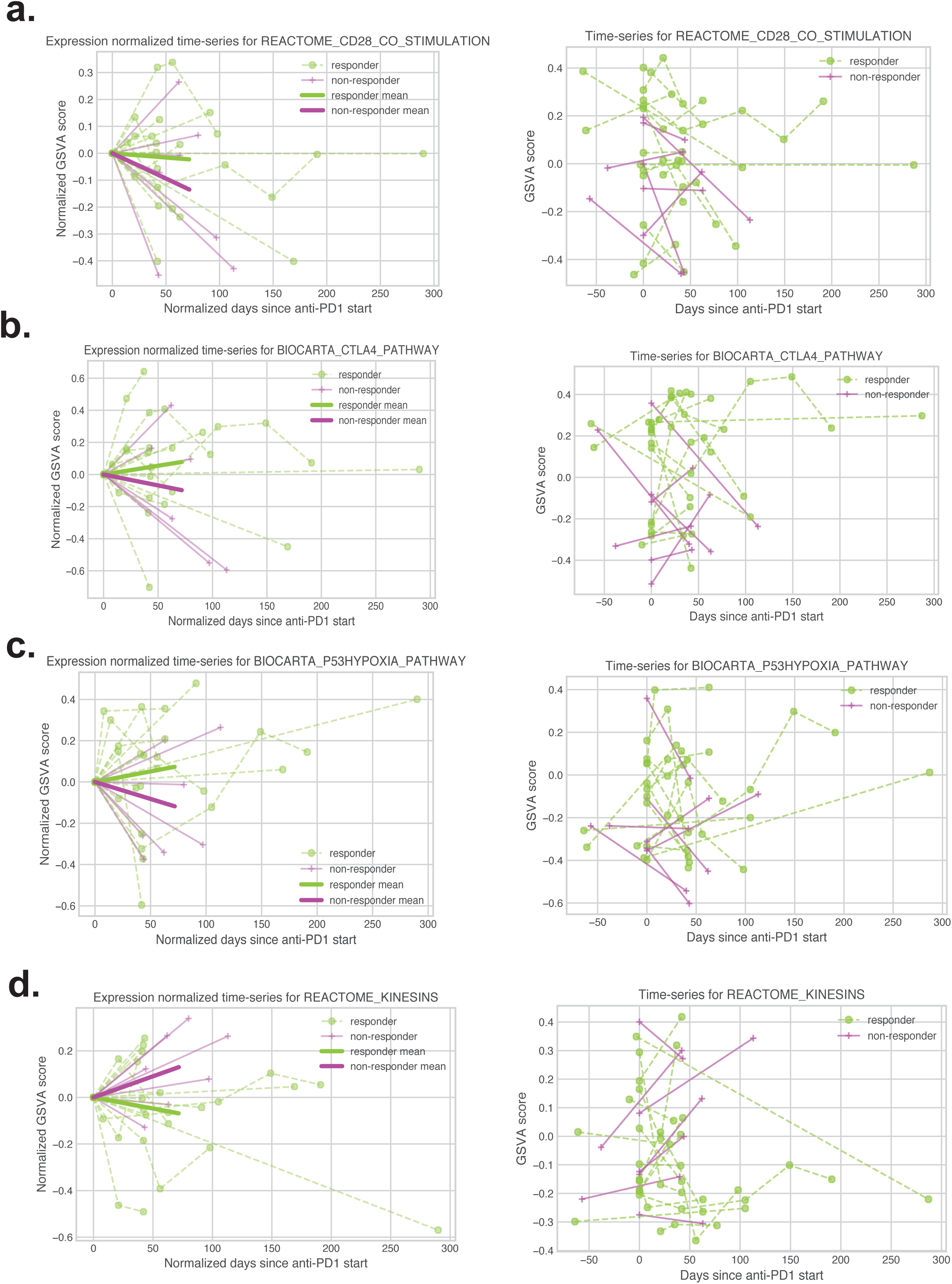
(a-d) Normalized and un-normalized time-series plots showing time dynamics for selected pathways. The plotting was performed per methodology previously discussed in captions for Fig. 2b, Fig. 2d and SFig 5.

**Supplementary figure 7:**
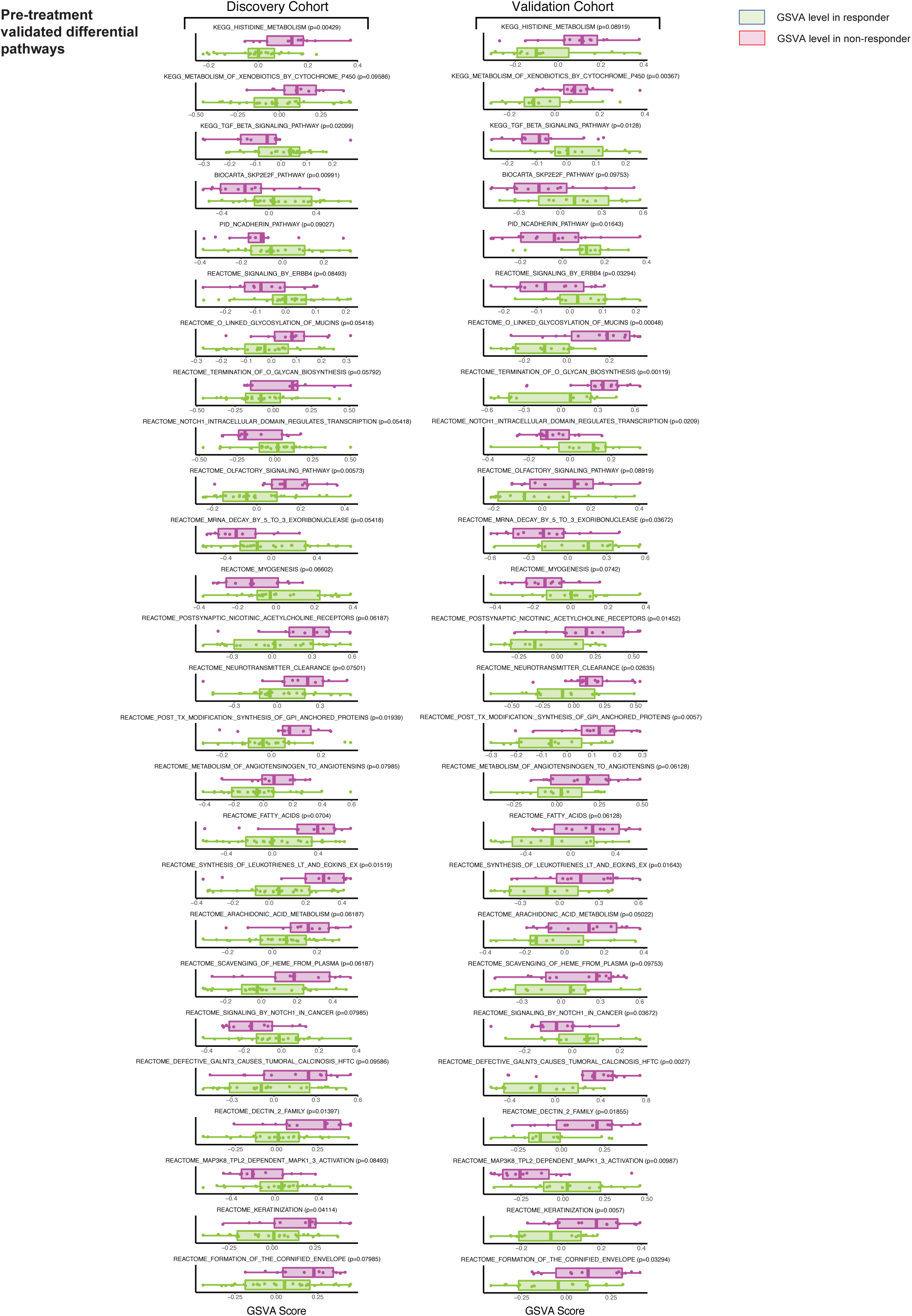
Kaplan-Meier progression-free and overall survival curves for selected genes that showed significant or near-significant differences between high-expressed and low-expressed patients.

**Supplementary figure 8:**
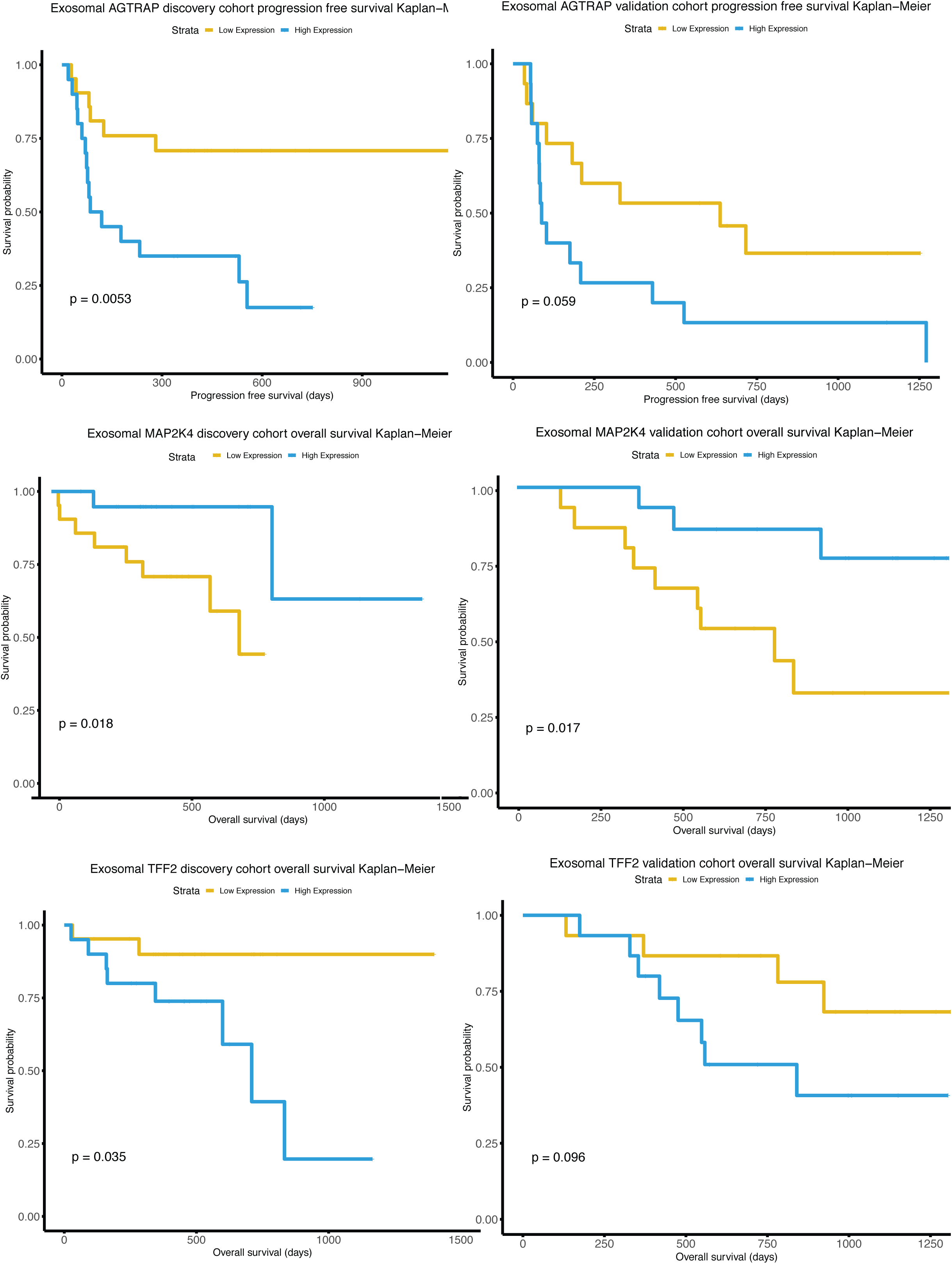
Kaplan-Meier progression-free and overall survival curves for selected genes that showed significant or near-significant differences between high-expressed and low-expressed patients.

**Supplementary figure 9:**
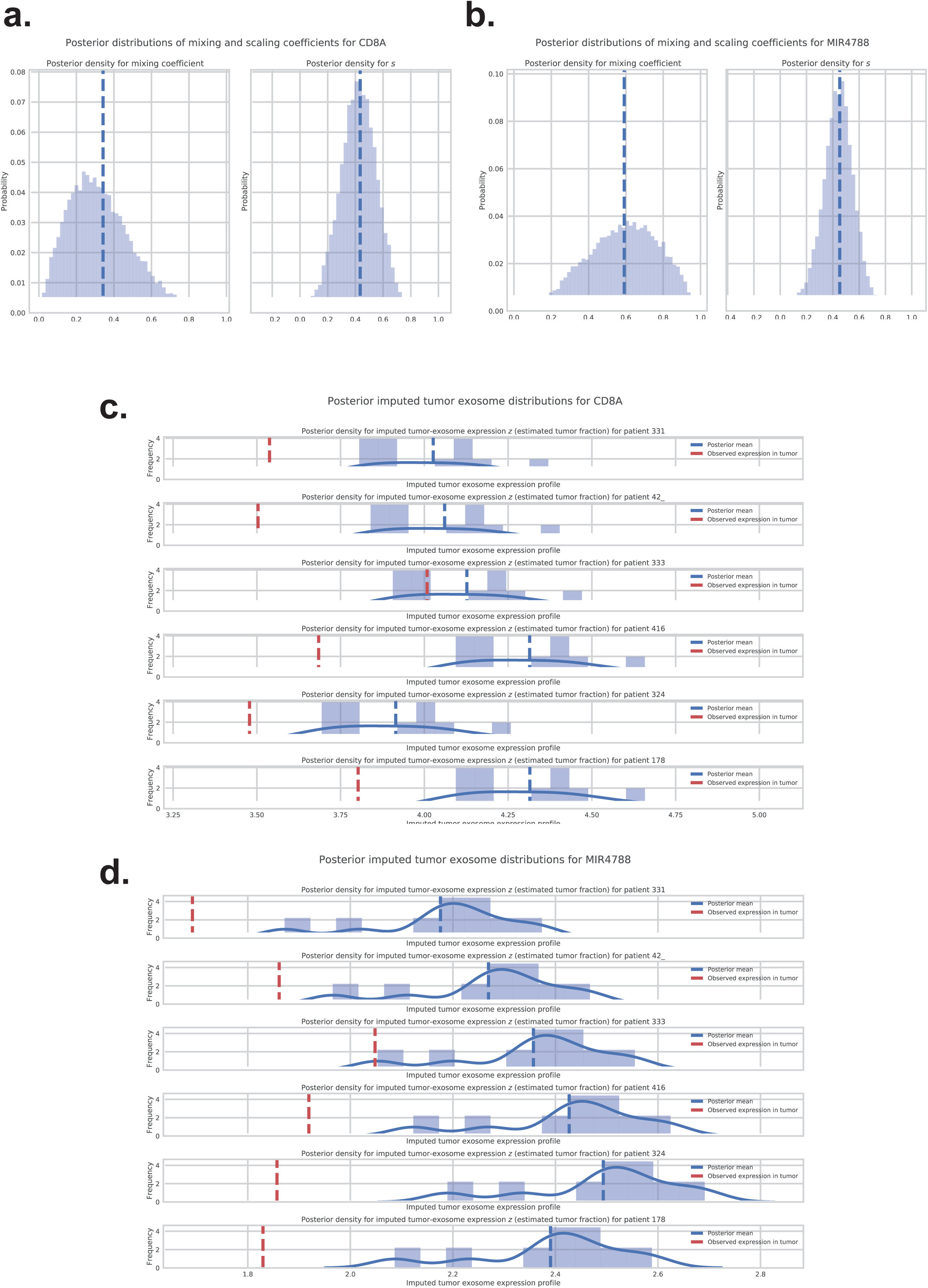
Example of per-patient imputed tumor-exosome expression from our Bayesian deconvolution model. (a-b) Posterior estimates for two illustrative genes for high immune fraction (CD8A) and high tumor fraction (MIR47888). (c-d) Predicted tumor-derived exosomal expression from our deconvolution model for CD8A and MIR4788 for a subset of patients.

## Supplementary Tables

**Supplementary Table 1:** Worksheet 1: Anonymized metadata of patient cohort from discovery cohort. Worksheet 2: anonymized metadata of patient cohort from validation cohort. Additional columns in worksheet 2 denote driver mutations detected by SNaPShot tumor sequencing and the specific mutations validated in our exoRNA-seq mutational analysis.

**Supplementary Table 2:** GO pathways enriched in exosome-unique or tumor-unique samples.

**Supplementary Table 3:** Validated DEGs and associated statistics from limma from our on-treatment DEG analysis.

**Supplementary Table 4:** Validated DEGs and associated statistics from limma from our pre-treatment DEG analysis.

**Supplementary Table 5:** Estimated mixing fractions from our exosomal deconvolution model running in multi-gene mode for all genes. Full details regarding the model are available in the supplemental note.

## Methods

### Tumor cell lines

Melanoma cell lines (A375, RPMI 7951, SK-MEL-30, SK-MEL-2, MeWo) were purchased directly from the American Type Culture Collection (ATCC) and maintained in culture per ATCC recommendations. The A375 cell line was maintained in Dulbecco’s minimal essential medium (DMEM), whereas RPMI 7951, SK-MEL-30, SK-MEL-2, and MeWo cell lines were cultured in RPMI-1640 media. All growth media consisted of media supplemented with 10% FBS, and 100 I.U./mL penicillin, 100 μg/ml streptomycin, and 0.292 mg/mL L-glutamine. Cells were grown on plates and incubated at 37°C with a humidified atmosphere of 5% CO_2_ in air.

### Patient samples and plasma isolation

Serial tumor and blood samples were collected from patients with melanoma under protocols approved by the Institutional Review Board at the Massachusetts General Hospital. Patient samples were linked to clinical data in a retrospective database. Blood was collected in sodium citrate cell preparation tubes, with plasma isolated after centrifugation at room temperature for 25-30 minutes at a relative centrifugal force of 1800. Plasma was then frozen and stored at −80°C until use. Peripheral blood mononuclear cells (PBMCs) are collected from the same sodium citrate tubes, washed in phosphate buffered saline, resuspended in DMSO with 90% FBS, slow frozen at 1°C per minute, and stored at −80°C until use.

### Isolation of exosomes

*Cell lines:* When 150mm plates reached between 50 – 70% confluence, depending on the doubling time of the cell line, the media was replaced with the appropriate media containing exosome-depleted FBS and the media was harvested after 48 hours. Exosomes were isolated from cell-conditioned media using serial centrifugation to removal cellular debris and filtration with a 0.8µM or smaller filter (Millipore) followed by ultracentrifugation as previously described ^32^. Briefly, cell-conditioned media was collected and centrifuged at 3,000 revolutions per minute (rpm) for 10 minutes at 4°C after which the supernatant was decanted and filtered using a 0.45µM or 0.8µM filter. The filtered supernatant then underwent ultracentrifugation at 150,000xg for 120 minutes. The pellet was then washed in PBS and underwent a second round of ultracentrifugation for 90 minutes. The exosomes were then resuspended in cold RPMI media on ice, and then frozen and stored at −80°C.

*Plasma:* All exosome for RNA transcriptomic analysis from plasma were isolated from 1ml of plasma using column isolation (Qiagen exoRNAeasy midi kit). Column isolation using the Qiagen exoRNeasy serum/plasma midi kit resulted in direct isolation of exosomal RNA (10–30ng RNA/ml). Approximately 1 in 10 patients had an additional 1ml of plasma isolated in parallel using ultracentrifugation (for quality control studies only, Fig. S2). For ultracentrifugation, plasma was thawed and filtered through a 0.2µM filter prior to ultracentrifugation as described for cell lines, above.

### Nanoparticle tracking analysis, electron microscopy, Western Blot analysis

*Nanoparticle tracking analysis:* Nanoparticle tracking analysis (NTA) using the Nanosight LM10 (Malvern) was employed in order to assess size distribution and concentration of particles in cell cultures and selected patient samples. Samples were diluted in PBS either 1/500 or 1/1000 according to Nanosight instruction manual.

*Transmission electron microscopy:* Electron microscopy was used to confirm the presence of exosomes in cell culture and selected patient samples by size and morphology. Isolated exosome suspensions were diluted 2:1 in 1xPBS and 8-10µl aliquots of each diluted sample were placed on formvar-carbon coated Ni mesh grids; samples were allowed to adsorb for 15 minutes. Following adsorption, grid preparations were placed on drops of primary antibody CD9, rabbit monoclonal (D801A), Cell Signaling #13174, diluted 1:25 (dilutions made in DAKO antibody diluent). Samples were allowed to incubate in primary antibody for at least 1 hour at room temperature, then rinsed on drops of PBS and incubated in drops of a secondary gold conjugate at least 1 hour at room temperature: Goat anti-rabbit 10nm IgG (Ted Pella #15726). Grids were then rinsed on drops of 1xPBS, then distilled water, contrast-stained for 10 minutes in droplets of chilled tylose/uranyl acetate, and air-dried prior to examining in a JEOL JEM 1011 transmission electron microscope at 80 kV. Images were collected using an AMT digital camera and imaging system with proprietary image capture software (Advanced Microscopy Techniques, Danvers, MA).

*Protein isolation from cell lines or exosomes:* protein was isolated using RIPA buffer supplemented with protease inhibitors.

*Protein Quantification:* Protein concentration of exosomal samples were determined using the *DC*™ Protein Assay (Bio-Rad) according to the manufacturer’s protocol.

*Western Blot*: Western blot was performed on samples of exosomes isolated from cell lines (Fig. S2) to confirm their presence in the samples. Antibodies for exosomal markers CD9^33^ were used as a positive marker given their consistency in expression across exosomal samples, whereas calnexin, an endoplasmic reticulum protein, is not found in exosomes and was used as a negative control. Actin was used as a loading control for cell lines.

### RNA extraction

#### Cell lines

Cells were trypsinized, washed in media for trypsin deactivation, then washed in PBS, counted, and pelleted. The cell pellet was resuspended in Trizol at a concentration of 5 x 10^6^ cells per 1mL and RNA isolated according to manufacturer’s instructions. RNA was then quantified using a NanoDrop spectrometer.

#### Cell line derived exosomes

RNA was extracted from exosomes using Qiagen exoRNeasy kit.

#### Tumor

Formalin-fixed tissue was analyzed to confirm that viable tumor was present via hematoxylin and eosin (H&E) staining. DNA was extracted from snap frozen tissue using Qiagen’s AllPrep DNA/RNA FFPE Kit.

#### Peripheral blood derived/plasma exosomes

Total exosomal RNA was extracted from thawed patient plasma using the Qiagen exoRNeasy Serum/Plasma Midi Kit as per the manufacturer’s protocol. Briefly, 1mL of plasma was thawed and filtered using a 0.8µM or smaller filter (Millipore). The sample was then mixed with binding buffer and placed on a spin column. After a wash step, the exosomes were lysed on the column using QIAzol and the eluate was then treated with chloroform to achieve phase separation. The aqueous phase was combined with 100% ethanol and then underwent column extraction with wash steps and final elution of exosomal RNA in RNase-free water.

### Discovery cohort microarray processing

We performed microarray analysis (via a subcontract to Thermo Fisher Scientific) utilizing Applied Biosystems GeneChip Human Transcriptome Array 2.0. Raw .cel files provided by ThermoFisher was read using the R package ‘oligo’. We performed background subtraction, quantile normalization, and summarization via the Robust Multichip Average (rma) algorithm using the R package ‘oligo’. We used the R package ‘pd.hta.2.0’ to provide functional annotations for the probes. We filtered all probes that did not map to an annotated gene, as well as duplicate probes. We corrected for batch effects using the ‘ComBat’ algorithm from the R ‘sva’ package^34^. For analyses that required validation by exoRNA-seq, the resulting matrix was further corrected for platform-specific effects using ‘ComBat’ and was quantile normalized (fully described in the RNA-seq processing section). Analyses that only utilized microarray data (SFig. 1b) was performed using the non-platform-corrected data, since cross-platform normalization with exoRNA-seq log2(TPM+1) may unduly bias microarray-only analyses.

### Validation cohort RNA-seq processing

Raw Illumina .fastq files were first filtered using ‘fastp’^35^ with default settings and then aligned to human hg38 transcriptomic reference using STAR v2.4.1 with default settings. To summarize the count data from the aligned .bam files, we utilized the function ‘featurecounts’ function from the R package ‘Rsubread’. Next, we derived log2(TPM+1) values using a custom R function. To make the RNA-seq count matrix co-analyzable with our microarray data, we essentially treated the resulting ComBat-corrected log2(TPM+1) matrix as an additional microarray batch. To remove platform-based effects, we concatenated log2(TPM+1) matrix with the microarray expression matrix to make a joint data matrix. We used the ‘ComBat’ algorithm from the R ‘sva’ package followed by quantile normalization using R ‘preprocessCore’ library on this joint data matrix. The resulting platform-normalized matrix was then split back into exoRNA-seq and microarray matrices for relevant downstream analyses.

### Differential expression analysis

To calculate differential expression in our microarray-based discovery and our exoRNA-seq validation cohort, we used the R ‘limma’ package to compute the p-values that corresponded to the comparisons. For the samples that had multiple replicates, we modeled biological replicates as a random effect. Confounding variables such as age and prior immunotherapy treatment were tested for association against ICI response and did not exhibit significant associations, and, as a result, they were not included as covariates. To find the top differentially expressed genes, we utilized limma’s Empirical Bayes linear modeling framework. Differentially expressed genes were defined by 1.5 log-fold change between responders and non-responders and a nominal p-value cutoff of p=0.1 from limma. In order to be considered validated, a gene has to be fulfill both the nominal p-value cutoff and log-fold change cutoff in both the validation and discovery cohorts. We note that this nominal p-value cutoff is ordinarily insufficient to control for false positive discoveries in a single cohort study; however, we require explicit confirmation for putative discovery cohort DEGs in our validation cohort, thus the combined false positive rate for a gene to be both falsely discovered and falsely validated is substantially lower than what the nominal p-value cutoff would suggest.

### Concordance and differential pathway analysis

Concordance analysis was performed by first binarizing the mean expression values of either the cell-line or patient data based on a 1.5 log expression cutoff using non-platform-corrected microarray data. If a gene is either present or absent in both groups, it is labeled as concordant. We averaged the expression signal from two pre-treatment patients present in the pretreatment time point; the post-treatment replicates were considered separately since they were from separate time points. To find the canonical (C2) MSigDB^13^ pathways that are significantly different between responders and non-responders in both the discovery and validation cohorts, we utilized the Gene Set Variation Analysis (GSVA)^14^ program with default settings to generate per-patient GSVA scores (a normalized statistic summarizing enrichment relative to the entire cohort analogous to ssGSEA scores) across our platform-corrected discovery and validation cohorts datasets. We then used a Mann-Whitney U-Test to test for differential GSVA scores between responders and non-responders. Similar to the rationale used for DEG analysis, we utilized a nominal cutoff p-value cutoff of 0.1 to flag differential pathways (Fig. 2a and Fig. 3a). A pathway was considered validated if it achieved significance in both the discovery and validation cohorts.

### Survival analysis & time-series analysis

To compute and plot the Kaplan-Meier curves, we utilized overall survival data and censoring information as inputs into the Kaplan-Meier computation and plotting functions in the R package ‘survminer’. To generate the time-series plots in Fig. 2b, we utilized the R Gene Set Variation Analysis (GSVA)^14^ package to generate a normalized enrichment score for each sample for target KEGG pathways. To generate the per-patient time dynamic plots illustrated in Fig 2c, we normalized discovery cohort expression data

### Building a predictive classifier

To build a predictive classifier from our selected pre-treatment DEGs in our discovery cohort, we first divided the patients into K=5 distinct cross-validation sets using the function ‘StratifiedKFold’ from the python library ‘sklearn.model_selection’. For each fold, an random-forest classifier was trained using the python library ‘sklearn’^31^, the combined predictions across all K=5 folds for three different models with varying number (10, 20, 30) of trees are plotted in Fig 3c. The features used were the validated pre-treatment DEGs presented in Fig. 3c and STable 4. Next, using the hyperparameters from the best-performing discovery cohort data, we next trained a random forest on the full dataset first plotted the internal cross-validation results, and performance was evaluated by calculating the area under the curve (AUC) of the receiver operating characteristic (ROC). We note that the model presented here is only intended to demonstrate the predictive power of validated pre-treatment DEGs on survival.

### Exosomal deconvolution modeling

We formulated a Bayesian probabilistic model to deconvolve for each gene the observed mixed plasma-exosomal expression profiles into two components: tumor and non-tumor. We explicitly model the enrichment/depletion of RNA abundance as a result of export from the tumor to tumor-derived exosomes by leveraging information from patient-derived exosomes. To model the non-tumor component, we utilize exosomal transcript abundances from healthy controls. A full mathematical description of the models and the source code are presented in the Supplementary Note. The model was implemented in the probabilistic programming language Stan^36^ through its python interface. Inference for key genes were conducted using the No-U-Turn Hamiltonian Monte Carlo sampler^37^. We performed 10,000 iterations with 4 chains in order to generate random samples; all other parameters were set using the pystan defaults. Semi-informative priors for several variables of interest were selected due to the limited availability of the data; we utilized a Beta(2,2) prior and a Normal(0,2) prior for the scaling coefficient for the mixing fraction to (1) model our prior belief regarding the predicted nature of mixing & scaling and (2) to constrain the posterior results somewhat in cases where certain genes with low expression but high variability in measurements may yield unreasonable mixing fraction results. We used a weakly informative prior on the variance parameter. Our model is able to return full posteriors for: the “packaging” coefficient, mixing fraction, and each patient’s unobserved tumor profile (Figure S7). Although our full probabilistic algorithm returns the full posteriors for many parameters of interest, there is a significant computational cost associated with running MCMC on tens of thousands of genes. Thus, we also created a simplified version of our model that is far more computationally efficient but only returns maximum a posterori (MAP) estimate of the mixing fraction instead of giving full posteriors estimates of several parameters. It does so by utilizing the Sequential Least SQuares Programming (SLSQP) algorithm from the python ‘scipy.optimize’. We utilized this simplified model to generate the full list of mixing fraction in Fig. 4b-f and Table S5.

### Mutational calling and analysis from exoRNA-seq data

To derive the mutational information shown in Table S1 and Fig. 4g-h, we first mapped all reads using ‘bwa’ mem v0.7.17 (with default settings) against reference human hg38 reference; ‘bwa’ was chosen to include reads lying outside of the reference transcriptome with potentially useful mutational information. Next, we used GATK ‘HaplotypeCaller’ submodule (with default settings) to call mutations from our patient RNA-seq libraries against hg38 reference and compared the resulting mutational information with summaries of SNaPshot panel sequencing results from clinical records. SNaPshot is a multiplexed PCR assay aimed identifying somatic variants in 70 different loci from 15 cancer genes; the SNaPshot assay was performed by MGH’s pathology department. We utilized a custom python script to overlap the resulting .vcf files produced by GATK against the ‘CosmicCodingMuts.vcf’ file downloaded from the COSMIC mutation database v89.

## Acknowledgments

The authors would like to thank Dr. Liang He, Dr. Yongjin Park, Dr. Yue Li, and Dr. David Liu for statistical consultations. We would like to thank Dr. David Ting and members of his lab for allowing us to perform exoRNA-seq in their laboratory.

## Contributions

G.M.B, G.G.K, S.C. designed the experiment. W.A.M, J. C-G., S.C., D. P., M.D.M. carried out the exosome experiments. A.S. and I.C. performed the bioinformatic and statistical analysis. A.S. designed and implemented the deconvolution model. D. J. was a material contributor. A.S., A.M., K.T.F., R.J.S, M.K., and G.M.B were involved in data interpretation and integration. A.S., G.M.B, J. C-G., G.G.K, and M.K. wrote the manuscript.

## Data Availability

The raw microarray chip and exoRNA-seq files, processed data files, and metadata has been uploaded to NCBI’s GEO database (GSE Accession Number #, submission in progress).

## Code Availability

Stan code used for the single-gene deconvolution model and python code used for the multi-gene deconvolution model are provided in the Supplemental Note.

## Supplemental note: exosomal deconvolution model

In this supplemental note, we provide the mathematical background underlying our deconvolution model, as described in Figure 4a of “Plasma-derived exosomal analysis and deconvolution enables prediction and tracking of melanoma checkpoint blockade response”. The rationale for the model is introduced in the main text. To summarize, we want to infer the contribution of the tumor-derived exosomal component and non-tumor derived (interchangeably referred to as “immune” and “non-tumor”) component to the observed plasma-derived exosomal transcriptomic profile. In contrast to exisiting deconvolution models designed for bulk deconvolution (e.g., CIBERSORT[1]), our model explicitly models the changes in transcript abundance as a result of export/packaging from the transcript abundance in the tumor to the transcript abundance. All of the data shown both here and in the main-figures related to deconvolution utilized only non-platform-corrected discovery cohort microarray data, since the discovery cohort exoRNA-seq included only plasma-derived exosomal samples and thus were not suitable inputs for our deconvolution model (see Supplementary Figure 1b for more detailed information regarding our analysis methodology). We created two versions of the deconvolution probabilistic model. In **single-gene mode**, the probabilistic model is fully fitted using the No-U-Turn Sampler (NUTS) Hamiltonian Monte Carlo (HMC) algorithm[2] and full posterior estimates for all the relevant parameters are returned. This model is fitted using the probabilistic programming language Stan[3]. This mode is designed for in-depth analysis of a single gene (or few genes), or situations where inferred parameters (e.g., scaling parameter, patient inferred tumor-exosome expression) of interest requires a full posterior estimate. Due to the time and resource intensive nature of the fitting process, it is impractical to perform full MCMC inference when we want to analyze the deconvolution profiles for tens of thousands of genes. Thus, we included a second mode, a **multi-gene mode**, in which we fit a simplified version of the single-gene model using Scipy’s implementation sequential least squares programming (SLSQP) to return a point estimate of the mixing coefficient. The single-gene model not only returns posterior distribution mixing fraction, but also the full posterior distribution of the scaling coefficient, which allows per-patient imputation of the tumor-derived exosomal fractions (Supplemental Figure 9); however, the multi-gene model only returns a single maximum a posterori (MAP) estimate of the mixing fraction. We envision the usage of the single-gene model in cases when a specific gene needs to carefully dissected and more robust inference is required, whereas the multi-gene model can be used on large-scale transcriptomic datasets to infer population-wide mixing fractions.

### 1 Single-gene Bayesian probabilistic model

- *N* : number of patient derived tumor profiles and tumor exosomal profiles
- *M* : number of cell-line tumor and tumor exosomal profiles
- *x_i_*: the *i*th patient’s observed tumor expression for the current gene
- *y_i_*: the *i*th patient’s observed peripheral-blood derived expression for the current gene
- *w_j_*: the *j*th cell-line’s
- *α*: mixing fraction between tumor-component and immune-component
- 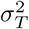: variance component
- *µ_I_* : Immune component mean (fixed parameter)
- 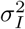: Immune variance component (fixed parameter)

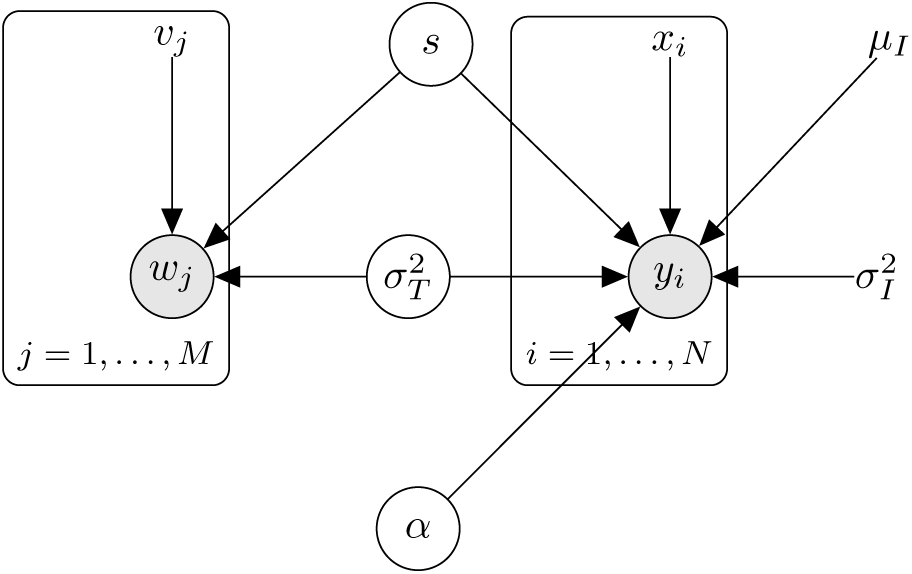

#### Prior specification

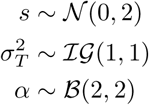

Where 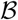 denotes the Beta distribution and 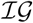 denotes the Inverse-Gamma distribution.

#### Data likelihood

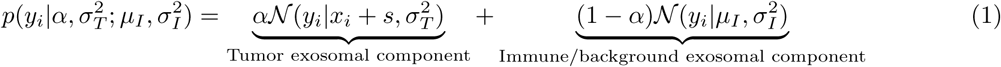

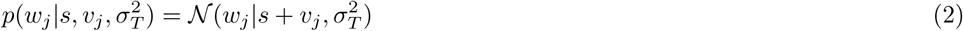

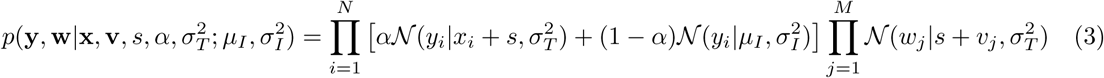

#### Full Posterior

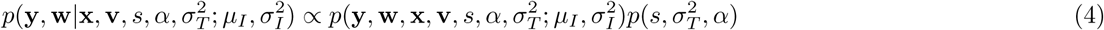

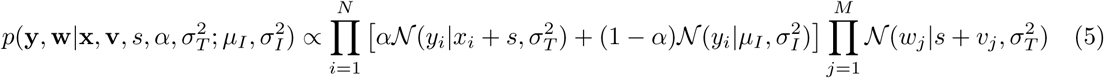

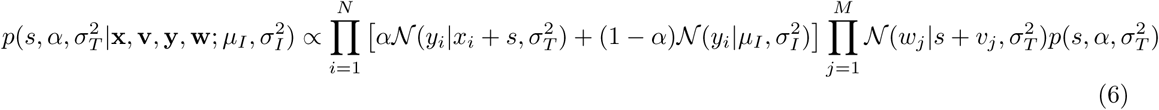

### 2 Multi-gene probabilistic model

For the multi-gene model, we utilize a simplified version of the model from the single gene model to reduce the time and computational intensiveness of the model. The multi-gene model only returns the mixing fraction α for the gene-of-interest, and as a result, it runs significantly faster and is able to process transcriptome-wide datasets on the order of hours instead of days on a single computer, unlike the more computationally intensive single-gene model.

The multi-gene model differs from the single gene model in several key respects:

- The inference of the scaling coefficient *s* is not done jointly with the inference of α and other parameters of interest. Instead, it is precomputed for each gene. To compute *s*, we take the mean difference between *v_j_* and *w_j_* for each gene and use this as a fixed constant.
- Thus, the inference of α is simplified into a much simpler problem that only depends on the right plate in the graph model depicted in section 1. Since we are interested in only in α, we optimize the log posterior formula using Sequential Least Squares Programming (SLSQP) from the python SciPy library with the appropriate constraints for α. The 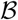(2, 2) prior for α is retained.

### 3 Validation of our deconvolution model using CIBERSORTx

**Figure S1:**
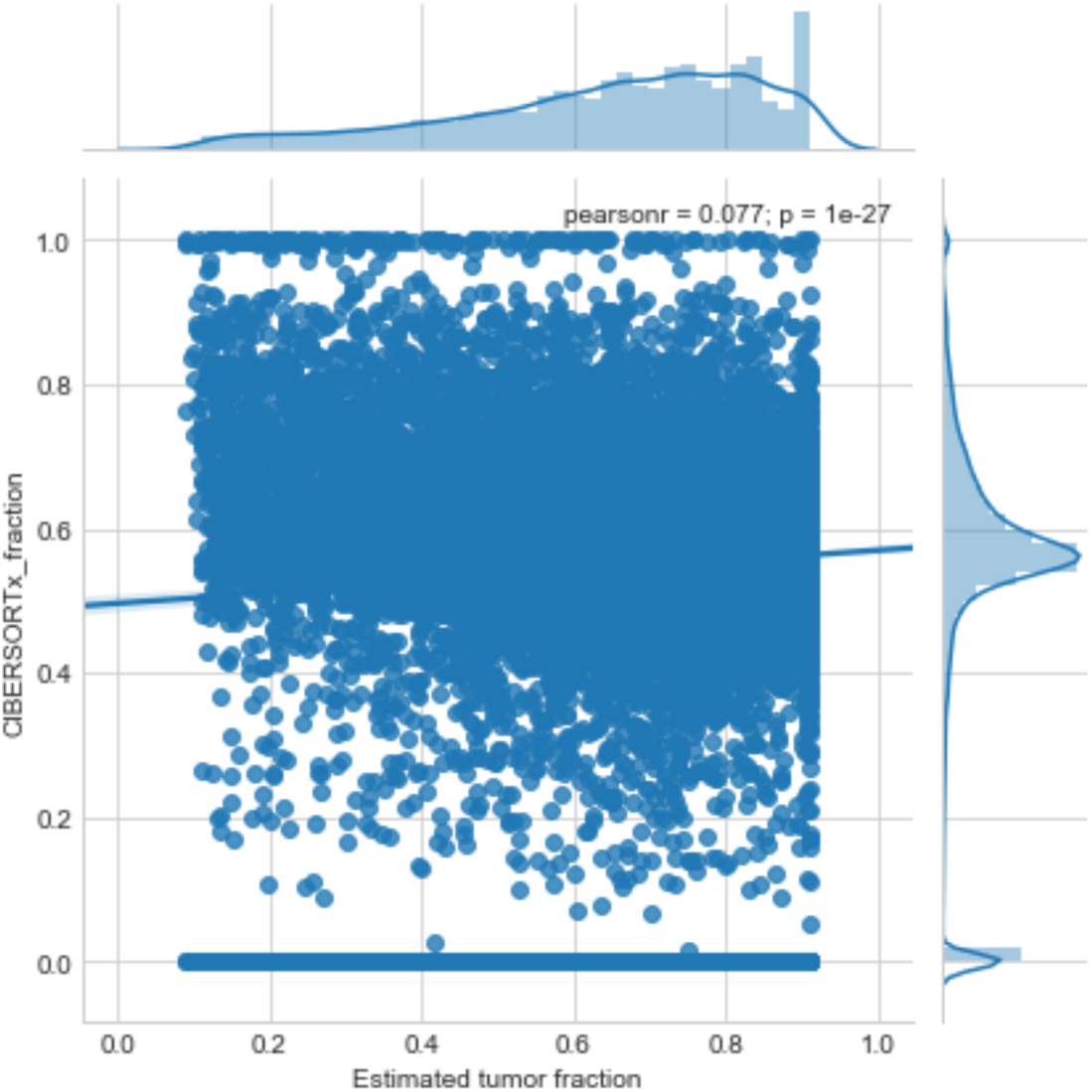
Scatterplot with line-of-best-fit and correlation between inferred CIBERSORTx and our deconvolution model inferred tumor fraction estimates on discovery cohort data

In order to provide evidence that our model is correctly partitioning genes into tumor and non-tumor components, we utilized CIBERSORTx[4]-a recently published bulk deconvolution program from Newman et al. that attempts to separate bulk transcriptomic profiles into component cell-type-specific profiles. In order to run the program, we utilized first inputted our discovery cohort (non-cross-platform corrected) microarray data matrix into the online CIBERSORTx web portal and utilized the melanoma reference profiles from Tirosh et al.’s *Science* 2016 single-cell dissection of metastatic melanoma that is provided by CIBERSORTx’s default online profiles. Since the algorithm generates estimates for all component cell-types instead of tumor-only profiles, we averaged the non-tumor cell-types in order to make it comparable to our non-tumor component estimation. A direct comparison of our values can be found in Supplemental Note Figure 1. We see that there’s a slight but significant correlation between CIBER-SORTx inferred-tumor fraction and tumor fraction inferred from our deconvolution model; however, it is clear from the CIBERSORTx density plot (y-axis) that the inferred tumor fraction is roughly normally distributed, an assumption that our model does not make (see density plot on x-axis). This continuous coding of tumor fractions hinders direct comparison of model predictions between CIBERSORTx and our model; thus, in order to better compare our cell-type predictions, we binarized model predictions for each gene as either tumor-derived or non-tumor derived using a cut-off of 0.5 as the threshold between tumor and non-tumor (same threshold used in the main manuscript). Using this cutoff, we can generate the confusion matrix found in Supplementary Note Table 1.

**Table 1:** Confusion matrix between CIBERSORTx and our deconvolution model using 0.5 tumor fraction as a cutoff between tumor and non-tumor binary classification of genes

|  | CIBERSORTx tumor | CIBERSORTx non-tumor |
| --- | --- | --- |
| Exosomal deconvolution tumor | 12984 | 2319 |
| Exosomal deconvolution non-tumor | 3719 | 1244 |

We can assess the concordance between binary predictions generated by CIBERSORTx and our model using values from the confusion matrix. This corresponded to the following binary classification statistics shown in Supplementary Note Table 2, using CIBERSORTx tumor predictions as “ground” truth and our deconvolution model estimates as predictions. We see that overall accuracy (0.70), sensitivity (0.78), precision (0.85), and F1-score (0.81) all support the ability of our deconvolution model to properly classify CIBERSORTx predicted tumor-derived genes. However, the two models’ predictions diverge significantly in terms of specificity (0.35) and negative predictive value (0.25), suggesting that the two models differ significantly in the overall prediction of non-tumor derived genes, with our model predicting a higher fraction of tumor-derived genes relative to CIBERSORTx. This is likely a result of the different distributional assumptions regarding tumor vs. non-tumor distributions between the two models (see Supplementary Note Figure 1). We reason that our model is likely to approximate reality more closely, based on known literature regarding significant increases in both overall and tumor-derived exosomal load in plasma during progression[5]. Furthermore, as mentioned in the main text, our model explicitly accounts for exosomal-specific characteristics - such as the differential exosomal packaging process - that bulk deconvolution techniques like CIBERSORTx does not account for. Additionally, our the underlying reference profiles is trained directly or inferred utilizing exosomal data, which is likely a far better approximation of the underlying mixture profiles in the context of cell-type deconvolution than bulk references. Though *in silico* independent validation via CIBERSORTx provides evidence for the validity of our deconvolution model predictions, particularly the prediction of tumor-derived genes, *in vivo* experimental evidence gathered via tumor vs. non-tumor derived exosomal selection/enrichment remains the gold standard to validate our model.

**Table 2:** Binary classification performance metrics generated from the confusion matrix in Table 1

| Metrics | Value |
| --- | --- |
| Accuracy | 0.70 |
| Sensitivity | 0.78 |
| Specificity | 0.35 |
| Precision | 0.85 |
| Negative Predictive Value | 0.25 |
| False Positive Rate | 0.65 |
| F1 Score | 0.81 |

### 4 Source Code

#### 4.1 Stan code for the single-gene model

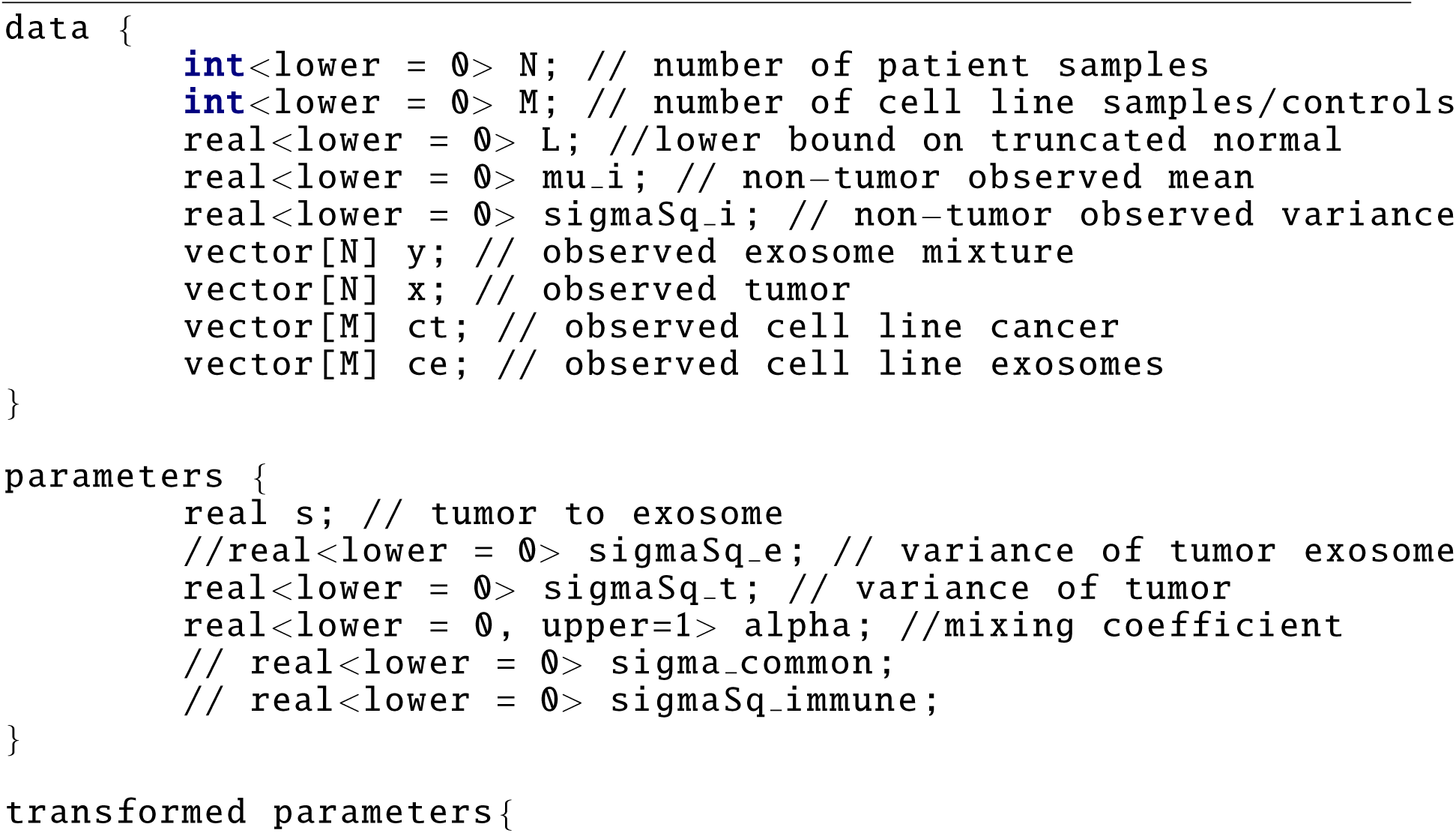
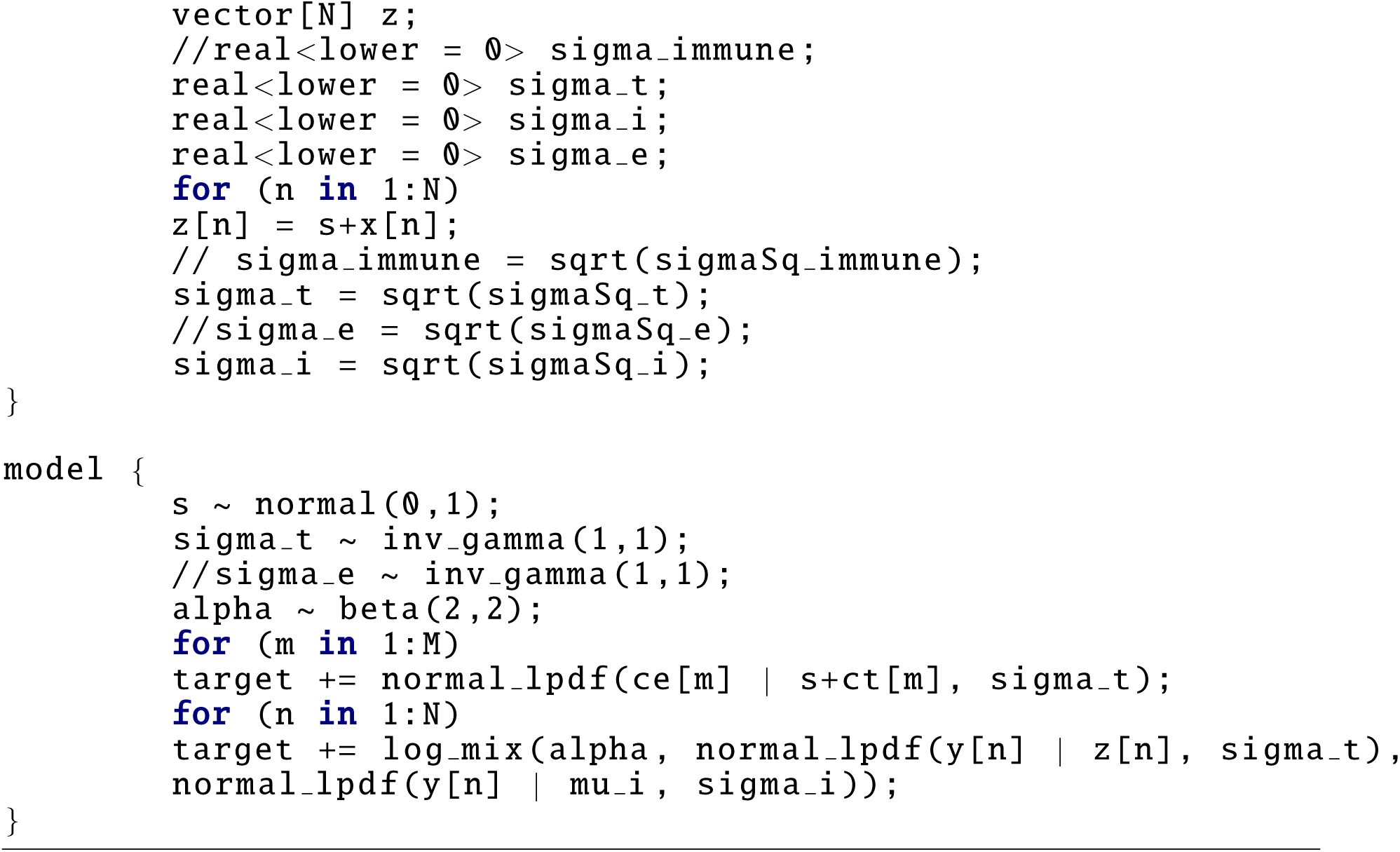

Inference was performed using pystan’s MCMC module with 10000 iterations and 4 chains, and all other parameters at default. The default pystan inference algorithm is the No-U-Turn Hamiltonian Monte Carlo (NUTS-HMC) sampler.

#### 4.2 Python inference code for the simplified multi-gene model MAP inference procedure

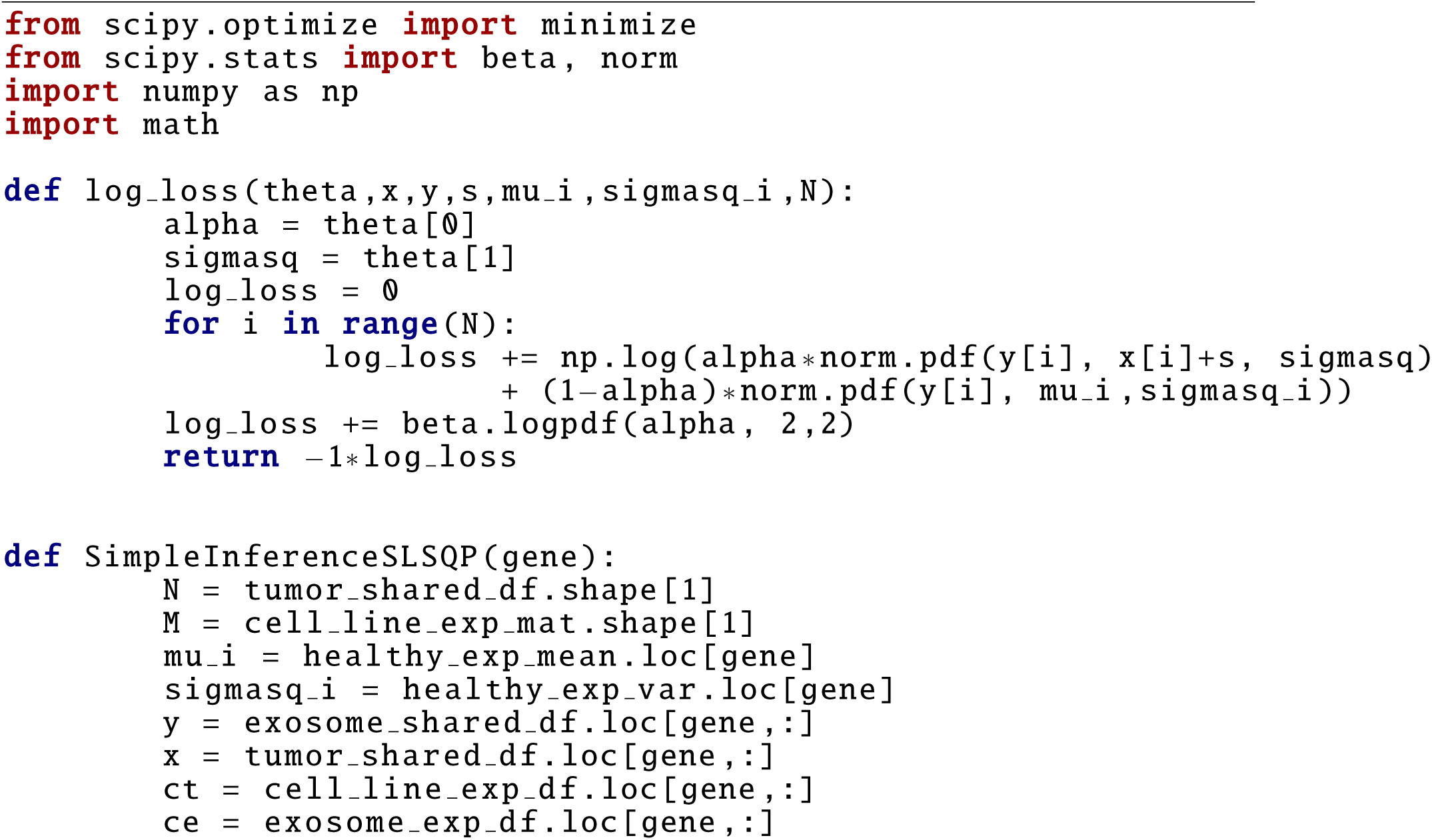
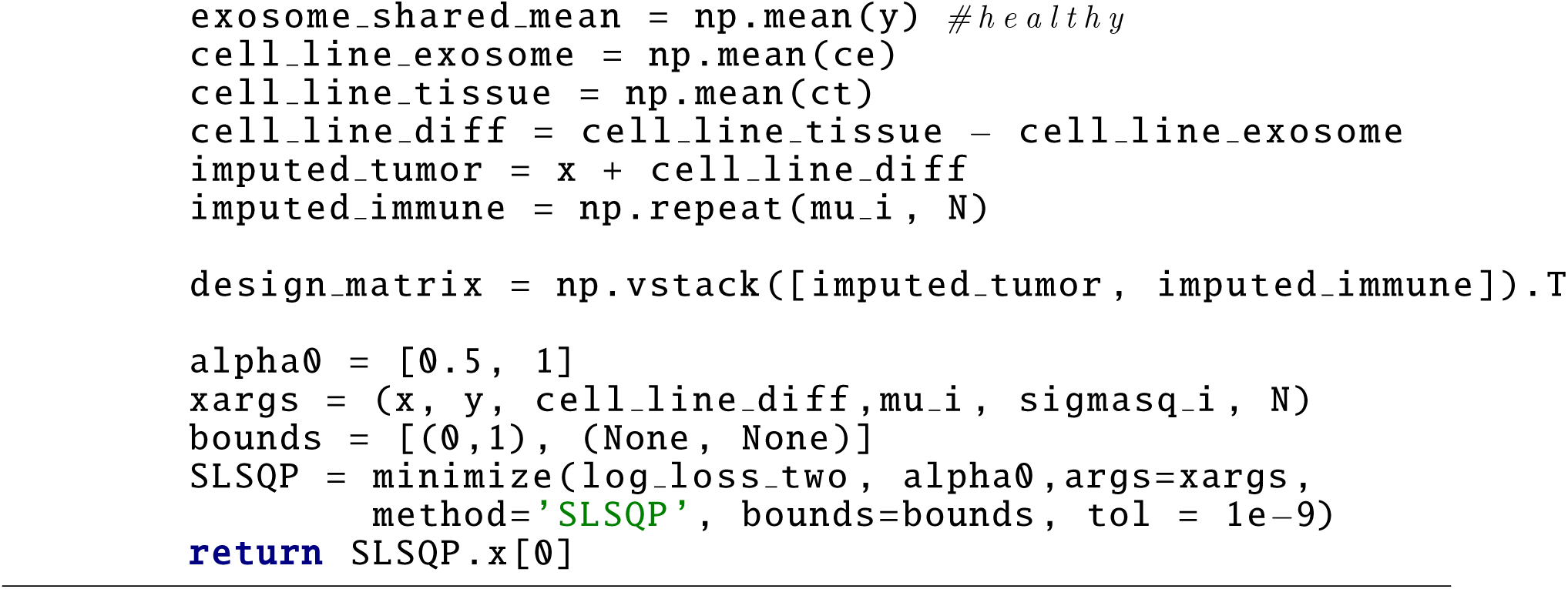

MAP inference was performed using the SLSQP package from the Python scipy.optimize library using default parameters. Note that the MAP estimates for α can and often differ from the single-gene probabilistic model - this is primarily due to the (1) the simplification of the probabilistic model for computational ease and (2) less robust/advanced inference algorithm relative to the NUTS-HMC used in the single-gene model.

## References

1. Hong, X. et al. Molecular signatures of circulating melanoma cells for monitoring early response to immune checkpoint therapy. Proc. Natl. Acad. Sci. U. S. A. 115, 2467–2472 (2018).

2. Ashida, A., Sakaizawa, K., Uhara, H. & Okuyama, R. Circulating Tumour DNA for Monitoring Treatment Response to Anti-PD-1 Immunotherapy in Melanoma Patients. Acta Derm. Venereol. 97, 1212–1218 (2017).

3. Riaz, N. et al. Tumor and Microenvironment Evolution during Immunotherapy with Nivolumab. Cell 171, 934–949.e15 (2017).

4. Alipoor, S. D. et al. The Potential Biomarkers and Immunological Effects of Tumor-Derived Exosomes in Lung Cancer. Front. Immunol. 9, 819 (2018).

5. Muller, L. et al. Exosomes isolated from plasma of glioma patients enrolled in a vaccination trial reflect antitumor immune activity and might predict survival. Oncoimmunology 4, e1008347 (2015).

6. McDonald, M. K. et al. Functional significance of macrophage-derived exosomes in inflammation and pain. Pain 155, 1527–1539 (2014).

7. Cai, Z. et al. Activated T cell exosomes promote tumor invasion via Fas signaling pathway. J. Immunol. 188, 5954–5961 (2012).

8. Greening, D. W., Gopal, S. K., Xu, R., Simpson, R. J. & Chen, W. Exosomes and their roles in immune regulation and cancer. Semin. Cell Dev. Biol. 40, 72–81 (2015).

9. Chatila, T. A. & Williams, C. B. Regulatory T cells: exosomes deliver tolerance. Immunity 41, 3–5 (2014).

10. Wong, K. H. K. et al. Whole blood stabilization for the microfluidic isolation and molecular characterization of circulating tumor cells. Nat. Commun. 8, 1733 (2017).

11. Newman, A. M. et al. Robust enumeration of cell subsets from tissue expression profiles. Nat. Methods 12, 453–457 (2015).

12. Roh, W. et al. Integrated molecular analysis of tumor biopsies on sequential CTLA-4 and PD-1 blockade reveals markers of response and resistance. Sci. Transl. Med. 9, (2017).

13. Liberzon, A. et al. Molecular signatures database (MSigDB) 3.0. Bioinformatics 27, 1739–1740 (2011).

14. Hänzelmann, S., Castelo, R. & Guinney, J. GSVA: gene set variation analysis for microarray and RNA-seq data. BMC Bioinformatics 14, 7 (2013).

15. Jenkins, R. W., Barbie, D. A. & Flaherty, K. T. Mechanisms of resistance to immune checkpoint inhibitors. Br. J. Cancer 118, 9 (2018).

16. Janghorban, M., Xin, L., Rosen, J. M. & Zhang, X. H.-F. Notch Signaling as a Regulator of the Tumor Immune Response: To Target or Not To Target? Front. Immunol. 9, 1649 (2018).

17. Sinnberg, T. et al. Wnt-signaling enhances neural crest migration of melanoma cells and induces an invasive phenotype. Mol. Cancer 17, 59 (2018).

18. Ni, X. et al. YAP Is Essential for Treg-Mediated Suppression of Antitumor Immunity. Cancer Discov. 8, 1026–1043 (2018).

19. Buchbinder, E. I. & Desai, A. CTLA-4 and PD-1 Pathways: Similarities, Differences, and Implications of Their Inhibition. Am. J. Clin. Oncol. 39, 98–106 (2016).

20. Memon, A. & Lee, W. K. KLF10 as a Tumor Suppressor Gene and Its TGF-β Signaling. Cancers 10, (2018).

21. Gattinoni, L. et al. Wnt signaling arrests effector T cell differentiation and generates CD8+ memory stem cells. Nat. Med. 15, 808–813 (2009).

22. Ohman Forslund, K. & Nordqvist, K. The melanoma antigen genes--any clues to their functions in normal tissues? Exp. Cell Res. 265, 185–194 (2001).

23. Luo, W., Friedman, M. S., Shedden, K., Hankenson, K. D. & Woolf, P. J. GAGE: generally applicable gene set enrichment for pathway analysis. BMC Bioinformatics 10, 161 (2009).

24. Hugo, W. et al. Genomic and Transcriptomic Features of Response to Anti-PD-1 Therapy in Metastatic Melanoma. Cell 165, 35–44 (2016).

25. Deng, J. et al. CDK4/6 Inhibition Augments Antitumor Immunity by Enhancing T-cell Activation. Cancer Discov. 8, 216–233 (2018).

26. Zhang, L. & Zhang, Z. Recharacterizing Tumor-Infiltrating Lymphocytes by Single-Cell RNA Sequencing. Cancer Immunol Res 7, 1040–1046 (2019).

27. Newman, A. M. et al. Determining cell type abundance and expression from bulk tissues with digital cytometry. Nat. Biotechnol. 37, 773–782 (2019).

28. Papadakis, K. A. et al. Krüppel-like factor KLF10 regulates transforming growth factor receptor II expression and TGF-β signaling in CD8+ T lymphocytes. Am. J. Physiol. Cell Physiol. 308, C362–71 (2015).

29. Goodman, A. M. et al. Tumor Mutational Burden as an Independent Predictor of Response to Immunotherapy in Diverse Cancers. Mol. Cancer Ther. 16, 2598–2608 (2017).

30. Tate, J. G. et al. COSMIC: the Catalogue Of Somatic Mutations In Cancer. Nucleic Acids Res. 47, D941–D947 (2019).

31. Pedregosa, F. et al. Scikit-learn: Machine Learning in Python. J. Mach. Learn. Res. 12, 2825–2830 (2011).

32. Théry, C., Amigorena, S., Raposo, G. & Clayton, A. Isolation and characterization of exosomes from cell culture supernatants and biological fluids. Curr. Protoc. Cell Biol. Chapter 3, Unit 3.22 (2006).

33. Lobb, R. J. et al. Optimized exosome isolation protocol for cell culture supernatant and human plasma. J Extracell Vesicles 4, 27031 (2015).

34. Johnson, W. E., Li, C. & Rabinovic, A. Adjusting batch effects in microarray expression data using empirical Bayes methods. Biostatistics 8, 118–127 (2007).

35. Chen, S., Zhou, Y., Chen, Y. & Gu, J. fastp: an ultra-fast all-in-one FASTQ preprocessor. Bioinformatics 34, i884–i890 (2018).

36. Gelman, A., Lee, D. & Guo, J. Stan: A Probabilistic Programming Language for Bayesian Inference and Optimization. J. Educ. Behav. Stat. 40, 530–543 (2015).

37. Hoffman, M. D. & Gelman, A. The No-U-Turn Sampler: Adaptively Setting Path Lengths in Hamiltonian Monte Carlo. arXiv [stat.CO] (2011).

## References

[1] Aaron M Newman et al. “Robust enumeration of cell subsets from tissue expression profiles”. In: Nature Methods 12.5 (Mar. 2015), pp. 453–457. issn: 1548-7091. doi: 10.1038/nmeth.3337. url: http://www.nature.com/doifinder/10.1038/nmeth.3337%7B%5C%%7D5Cnhttp://www.ncbi.nlm.nih.gov/pubmed/25822800%20http://www.nature.com/doifinder/10.1038/nmeth.3337.

[2] Matthew D. Hoffman and Andrew Gelman. “The No-U-Turn Sampler: Adaptively Setting Path Lengths in Hamiltonian Monte Carlo”. In: Journal of Machine Learning Research 15.April (2014), pp. 1593–1623. arXiv: 1111.4246. url: http://mc-stan.org/.

[3] A. Gelman, D. Lee, and J. Guo. “Stan: A Probabilistic Programming Language for Bayesian Inference and Optimization”. In: Journal of Educational and Behavioral Statistics 40.5 (2015), pp. 530– 543. issn: 1076-9986. doi: 10.3102/1076998615606113.

[4] Aaron M. Newman et al. “Determining cell type abundance and expression from bulk tissues with digital cytometry”. In: Nature Biotechnology 37.7 (July 2019), pp. 773–782. issn: 1087-0156. doi: 10.1038/s41587-019-0114-2. url: http://www.nature.com/articles/s41587-019-0114-2.

[5] Tao Huang and Chu-Xia Deng. “Current Progresses of Exosomes as Cancer Diagnostic and Prognostic Biomarkers”. In: International Journal of Biological Sciences 15.1 (2019), pp. 1–11. issn: 1449-2288. doi: 10.7150/ijbs.27796. url: http://www.ijbs.com/v15p0001.htm.

